# A familial natural short sleep mutation promotes healthy aging and extends lifespan in *Drosophila*

**DOI:** 10.1101/2023.04.25.538137

**Authors:** Pritika Pandey, P. Kerr Wall, Stephen R. Lopez, Olga S. Dubuisson, Elizabeth R.M. Zunica, Wagner S. Dantas, John P. Kirwan, Christopher L. Axelrod, Alyssa E. Johnson

## Abstract

Sleep loss typically imposes negative effects on animal health. However, humans with a rare genetic mutation in the *dec2* gene (*dec2^P384R^*) present an exception; these individuals sleep less without the usual effects associated with sleep deprivation. Thus, it has been suggested that the *dec2^P384R^* mutation activates compensatory mechanisms that allows these individuals to thrive with less sleep. To test this directly, we used a *Drosophila* model to study the effects of the *dec2^P384R^*mutation on animal health. Expression of human *dec2^P384R^* in fly sleep neurons was sufficient to mimic the short sleep phenotype and, remarkably, *dec2^P384R^* mutants lived significantly longer with improved health despite sleeping less. The improved physiological effects were enabled, in part, by enhanced mitochondrial fitness and upregulation of multiple stress response pathways. Moreover, we provide evidence that upregulation of pro-health pathways also contributes to the short sleep phenotype, and this phenomenon may extend to other pro-longevity models.

## Introduction

Sleep is an ancient behavior that is universally conserved among the animal kingdom (Keene and Duboue, 2018). However, a high degree of variability exists in the amount of time different species spend sleeping (Campbell and Tobler, 1984). Some species, such as *C. elegans*, only sleep during critical developmental transitions or injury (Raizen et al., 2008), while others, including many bat species, spend most of their life sleeping (Campbell and Tobler, 1984). Although work in the past few decades has led to a better understanding of the molecular mechanisms governing sleep homeostasis (Artiushin and Sehgal, 2017; Borb and Achermann, 1999; Vanini and Torterolo, 2021), why organisms require a certain amount of sleep is still a fundamental mystery.

In most species, it is evident that sleep is important for maintaining physiological health as inadequate sleep correlates with numerous health issues, such as hypertension, heart disease, metabolic disorders, cognitive impairment, neurodegenerative diseases, and even premature mortality (Cappuccio et al., 2011; Harper et al., 2001; Koh et al., 2008; Kwok et al., 2018; Laaboub et al., 2022; Mazzotti et al., 2014; Medori et al., 1992; Scullin and Bliwise, 2015; Sejbuk et al., 2022; Zhai et al., 2015). Moreover, a bidirectional relationship between aging and sleep exists; aging correlates with increased sleep disturbances, while reduced sleep accelerates aging phenotypes (De Nobrega and Lyons, 2020; Mattis and Sehgal, 2016). But, despite the strong link between sleep and maintaining cellular functions, there are rare examples of species that have adapted to cope with much less sleep compared to physiologically similar counterparts (Campbell and Tobler, 1984; Duboué et al., 2011). A striking example are populations of cavefish that have evolved to sleep up to 80% less but maintain a similar lifespan as their surface fish ancestors (Duboué et al., 2011). In recent years, natural short sleepers have even been identified in the human population that, despite sleeping less, do not exhibit adverse health issues that are typically associated with sleep deprivation. These examples hint that some organisms may have adapted reduced sleep requirements and that perhaps ectopically inducing pro-health mechanisms can influence the amount of daily sleep an organism needs. Understanding how these exceptional individuals compensate for less sleep may reveal unique strategies that can sustain health in sleep-deprived states as well as promote more general health.

One of the most well-studied examples of natural short sleepers in the human population are individuals with rare genetic mutations in the *dec2* gene (He et al., 2009). *Dec2* is a transcriptional repressor that, in mammals, is recruited to the *prepro-orexin* promoter and represses the expression of *orexin*, a neuropeptide that promotes wakefulness (Ganguly-Fitzgerald et al., 2006; He et al., 2009; Hirano et al., 2018). A single point mutation in *dec2* (*dec2^P384R^*) inhibits the ability of *Dec2* to bind the *prepro-orexin* promoter, resulting in increased *orexin* expression (Hirano et al., 2018). Consequently, wakefulness increases, and individuals sleep on average 6hrs/day instead of 8hrs/day (He et al., 2009; Hirano et al., 2018). Intriguingly, these natural short sleepers do not appear to exhibit any phenotypes typically associated with chronic sleep deprivation, and expression of the *dec2^P384R^* mutation in mice suppresses neurodegeneration (Dong et al., 2022; Gambetti et al., 1995; Pellegrino et al., 2014). Thus, it has been suggested that individuals harboring the *dec2^P384R^* mutation may employ compensatory mechanisms that allow them to thrive with chronic sleep loss. However, whether the *dec2^P384R^* mutation directly confers global health benefits has not yet been tested experimentally in any system.

In this study, we used a *Drosophila* model to understand the role of the *dec2^P384R^* mutation on animal health and elucidate the mechanisms driving these physiological changes. We found that the expression of the mammalian *dec2^P384R^* transgene in fly sleep neurons was sufficient to mimic the short sleep phenotype observed in mammals. Remarkably, *dec2^P384R^*mutants lived significantly longer with improved health despite sleeping less. In particular, *dec2^P384R^* mutants were more stress resistant and displayed improved mitochondrial fitness in flight muscles. Differential gene expression analyses further revealed several altered transcriptional pathways related to stress response, including detoxification and xenobiotic stress pathways, that we demonstrate collectively contribute to the increased lifespan and improved health of *dec2^P384R^* mutants. Finally, we provide evidence that the short sleep phenotype observed in *dec2^P384R^* mutants may be a result of their improved health rather than altered core sleep programs. Taken together, our results highlight the *dec2^P384R^* mutation as a novel pro-longevity factor and suggest a link between pro-health pathways and reduced sleep pressure.

## Methods

### Fly strains and rearing conditions

Fly stocks were raised at 25°C with 12:12h L:D cycle and fed on a standard cornmeal molasses medium. The following fly strains were obtained from the Bloomington *Drosophila* Stock Center (BDSC): *GR23E10-gal4* (49032), *UAS-dec2^WT^* (64227), *UAS-dec2^P384R^* (64228), *AMPKα* (32108), *UASp-foxo* (42221), *col4a1-gal4* (7011), and *elav-gal4* (8760). The following RNAi lines were obtained from Vienna *Drosophila* Resource Center (VDRC): *mtnB^i^* (106118), *mtnC^i^* (35816), *mtnD^i^*(330619), *CG11699^i^* (101491), *nmdmc^i^* (110198).*<ι><ιτ></i>*

### Generating LexAop-dec2 transgenic strains

The human *dec2* genes were PCR-amplified from *dec2^WT^ and dec2^P384R^* transgenic strains using the following primers:5’-GGGGACAACTTTGTATACAAAAGTTGTAATGGACGAAGGAATTCCTCATTTGC-3’ and 5’-GGGGACCACTTTGTACAAGAAAGCTGGGTATCAGGGAGCTTCCTTTCCTGGCTGC-3’. *dec2^WT^ and dec2^P384R^* transgenes were subsequently cloned into the pDONR P5-P2 Gateway vector (Invitrogen) using BP clonase (ThermoFisher Scientific, Cat# 11789020), to generate pENTR L5-*dec2^WT^*-L2 and pENTR L5-*dec2^P384R^*-L2, and the inserts were validated via DNA sequencing. pENTR L5-*dec2^WT^*-L2 and pENTR L5-*dec2^P384R^*-L2 plasmids were then individually combined with pENTR L1-13XLexAop2-R5 (Addgene #41433; (Petersen and Stowers, 2011)) and destination vector pDESTsvaw (Addgene #32318; (Petersen and Stowers, 2011)) using LR clonase (ThermoFisher Scientific, cat#12538120). The pDESTsvaw vectors contains a mini-white rescue gene to enable transgenic selection and an attB site to enable *PhiC31*-mediated site-specific integration. Injection services of Genetivision (Houston, TX) were used to insert transgenes at the vk27: (3r) 89e11 site using *phiC31*-mediated insertion. Transgenic lines were maintained over the 3^rd^ chromosome balancer TM6B using standard genetic crossing procedures.

### Sleep Analyses

Sleep analyses were performed using a *Drosophila* Activity Monitoring System (DAMS) from Trikinetics (Waltham, MA). In flies, sleep is defined as a quiescent period of five mins or longer (Hendricks et al., 2000). Male flies (7-10 days old) were loaded in 5 × 65 mm glass tubes with food on one side and were allowed to acclimate for approximately 24 hrs. Baseline sleep was measured as bouts of 5 min of rest and was recorded for 5 days. During the analysis, flies were subjected to a 12:12h L:D cycle in an incubator at 25° C. Sleep data was analyzed using ShinyR-DAM software (Cichewicz and Hirsh, 2018). For sleep rebound, baseline sleep was recorded for 24h before flies were subjected to 24h of sleep deprivation using a sleep nullifying apparatus (Melnattur et al., 2020), which tilts asymmetrically from −60° to +60° angle to mechanically displace flies 10 times per min. Rebound sleep was then recorded for 24h post-deprivation.

### Lifespan Assays

Adult *Drosophila* flies were collected within 24h of eclosion, transferred to fresh vials, and allowed to mate for 2-3 days to reach sexual maturity. Male flies were then isolated and transferred into fresh vials (20 flies per vial for a total of 6 vials). Lifespan experiments were conducted in an incubator with a controlled temperature of 25°C and a 12:12h L:D cycle. Flies were transferred into fresh vials every two days and dead flies were scored at the time of transfer. For lifespans under various stress conditions, 20 mM Paraquat, 12 μM Tunicamycin or 500 µM Rotenone was added directly to the food to induce oxidative stress, endoplasmic reticulum (ER) stress, or mitochondrial stress, respectively. Lifespans under sleep deprivation stress was performed by using a Sleep Nullifying Apparatus (SNAP) (Melnattur et al., 2020). Statistical analyses were performed using OASIS software (Han et al., 2016).

### Memory assay

To test memory function, we used an Aversive Phototaxis Suppression Assay. This assay is based on the principle that flies are naturally attracted to light, except when an aversive odor (Quinine hydrochloride dihydrate, MP Biomedicals) is simultaneously present. For each experiment, ∼20 adult male flies were transferred to an empty vial and starved for 6h before conducting the experiment to promote active foraging during the experiment. Before each experimental trial, flies were tested to determine whether they were positively phototaxic under normal conditions; flies were acclimated to the dark chamber for 30s, and flies that failed to migrate towards the light chamber after 25s were considered non-phototaxic and were censored from the experiment. For the remaining flies that were phototaxic, a filter paper soaked with quinine solution was inserted into the light chamber and 12 training trials were conducted. For each trial, flies were allowed 60s to migrate towards the light chamber. Flies that migrated towards the light chamber within 60s were scored as “Fail” and flies that stayed in the dark chamber scored as “Pass”. Immediately after 12 training trials, five test trials were performed to test short-term memory. In the test trials, the light chamber contained filter paper soaked with water. Flies that migrated towards the light chamber within 10s were scored as “Fail” and flies that remained in the dark chamber were scored as “Pass”. For long-term memory, the same flies were kept in vials with food for 4-5h before conducting five more test trials. For each test trial, the average pass rate for the five test trials was calculated for each individual fly.

### RNA sequencing

For each replicate, ∼70 whole flies of one week old were collected at ZT3, the time where there was most significant difference in their daytime sleep were used for RNA extraction. RNA extraction was done using standard a TRIzol TM reagent protocol (Thermo Fisher Scientific, cat# 15596018). Subsequently, genomic DNA was removed using a GeneJet RNA-purification kit (Thermo Fisher Scientific, cat# K0702). The concentration of purified RNA was measured using a nanodrop and quality was assessed using a Bioanalyzer. For each genotype, three independent biological replicates were sequenced.

For Illumina sequencing, at least 50 ng/μl of purified RNA for each replicate was sent to Novogene (Sacramento, CA) for cDNA library preparation and Illumina sequencing (Illumina NovaSeq 6000). For Nanopore sequencing, 10 μg of total RNA were diluted in 100 μl of nuclease-free water to prepare mRNA. Poly(A) RNA was separated using NEXTflex Poly(A) Beads (BIOO Scientific cat # NOVA-512980). Resulting poly(A) RNA was eluted in nuclease-free water and stored at –80°C. The quality of mRNA was assessed using a Bioanalyzer. 200 ng of input polyA + RNA was used to prepare cDNA libraries using a Direct cDNA sequencing kit (SQK-DCS109) and these were prepared according to the Oxford Nanopore recommended protocol. cDNA libraries were sequenced on a MinION using R9.4 flow cells.

### Differential expression analyses

For the Nanopore reads, we mapped the reads to the reference genome using Minimap2 (Li, 2018) with arguments (p=80 and N=100) as described in (Soneson et al., 2019). We then used Salmon (Patro et al., 2017) to quantify gene expression in alignment-based mode. For both the Illumina and Nanopore data, differential expression analysis among the 3 conditions (three biological replicates per condition) were performed using the DESeq2 (Love et al., 2017) R package (1.20.0). DESeq2 provides statistical routines for determining differential expression in digital gene expression data using a model based on the negative binomial distribution. The resulting P-values were adjusted using the Benjamini and Hochberg’s approach for controlling the false discovery rate. Genes with an adjusted p-value ≤ 0.05 and fold-change ≥ 1.5 found by DESeq2 were assigned as differentially expressed. We further analyzed the differentially expressed genes with enrichR (Chen et al., 2013) to look for enriched gene sets (adjusted p-value <= 0.05) with respect to KEGG (Ogata et al., 1999) and Gene Ontology (Ashburner et al., 2000). The results from both the Illumina and Nanopore data were combined using the Flybase gene identifiers and the final summary files are provided as supplemental.

For the Illumina datasets, reads were mapped to a reference genome (r6.39) of *Drosophila melanogaster* (Gramates et al., 2022) using HISAT2 (Pertea et al., 2016). We then used featureCounts v1.5.0-p3 (Liao et al., 2014) to count the reads mapped to each gene and calculate FPKM.

### qPCR methods

RNA was extracted from whole *Drosophila* animals and converted to cDNA using iScript™ cDNA Synthesis Kit (Bio-Rad, cat#1708891). Primers for qPCR were designed using IDT PrimerQuest (Fig. S7). Three experimental replicates per strain were analyzed and Actin was used as a housekeeping gene. qPCR was conducted using PowerUp^TM^ SYBR^TM^ Green Master Mix (ThermoFisher Scientific). qPCR was performed on a QuantStudio 6 Real-Time PCR system. Data were analyzed using standard ΔCT method. The 2^-ýýCT^ method was used to estimate the relative changes in gene expression. Data were normalized to the WT control.

### Mitochondrial respiratory capacity

OXPHOS and ET capacity of flight muscle homogenates was determined by high-resolution respirometry (Oroboros O2k; Innsbruck, Austria) as described previously (Wall et al., 2021).

Briefly, flies (∼5/biological replicates) aged 7-8 days were sedated by cold exposure at 4°C for 7-10 min. While sedated, thorax muscle was isolated from surrounding tissue and placed into ice-cold biopsy preservation solution (Doerrier et al., 2018). Thoraxes were then blotted dry, weighed, and placed into an ice-cold Dounce homogenizer containing mitochondrial respiration medium (MiR05) (Doerrier et al., 2018). Samples were homogenized for 20-30 seconds (7-9 strokes) and brought up to total volume with MiR05 (0.4 mg/mL final). Tissue homogenates were then transferred into an oxygraph chamber, containing 2 ml of MiR05, oxygenated to 600 μM, the chamber closed, and respiration was allowed to stabilize. Oxidative phosphorylated (OXPHOS) and electron transfer capacity was determined using the following concentrations of substrates, uncouplers and inhibitors: malate (2 mM), pyruvate (2.5 mM), ADP (2.5 mM), proline (5 mM), succinate (10 mM), glycerol-3-phosphate (15 mM), tetramethyl-p-phenylenediamine (TMPD, 0.5 μM), ascorbate (2 mM), carbonylcyanide-p-trifluoromethoxyphenylhydrazone (FCCP, 0.5 μM increment), rotenone (3.75 μM), atpenin A5 (1 μM), antimycin A (2.5 μM) and sodium azide (200 mM). Outer membrane integrity was confirmed by exogenous cytochrome c (7.5 μM).

### Supercomplex formation

Mitochondrial respiratory chain super complexes were resolved by blue native polyacrylamide gel electrophoresis (BN-PAGE) as described previously (Jha et al., 2016). Briefly, flight muscles (∼50/biological replicates) were dissected, minced, and homogenized in ice cold isolation buffer. Following centrifugation, the mitochondrial pellet was resuspended in isolation buffer and stored at −80°C until time of assay. Pellets were resuspended in ddH_2_O containing sample buffer, digitonin (5%), and coomassie G-250. Samples were loaded into a 3-12% NativePAGE gel and resolved by electrophoresis. Gels were stained with colloidal blue, and bands were visualized using iBright™.

### FAD Activity

FAD activity was determined colorimetrically by commercially available enzymatic assay (Abcam, ab204710). Isolated thorax muscles (∼5/biological replicate) were deproteinated (Abcam, ab204708), homogenized in ice cold FAS assay buffer, and centrifuged at 4°C at 10,000 x g for 5 minutes to remove insoluble material. Data are expressed as nmols of activity per minute per mg protein.

## Results

### Expression of human dec2^P384R^ in Drosophila sleep neurons reduces sleep and sleep rebound

Many regulatory mechanisms of mammalian sleep, including diurnal sleep-wake cycles, sleep rebound, and circadian rhythms are conserved in *Drosophila* (Dissel, 2020; Hendricks et al., 2000; Konopka and Benzer, 1971; Shaw et al., 2000). These aspects, coupled with their relatively short lifespans and robust genetic toolkits, make *Drosophila* an excellent model system to study the relationship between sleep and age-related animal physiology. Previously, it has been demonstrated that over-expressing the human *dec2^P384R^* mutant transgene in the *Drosophila* mushroom body (MB) mimics the short sleep phenotype observed in humans with the mutation (He et al., 2009). Although the MB encompasses sleep-promoting neurons, it also includes additional neurons not related to sleep regulation (Modi et al., 2020). Since this study, *gal4* drivers that express more specifically in sleep neurons have been developed and characterized, including GR23E10-*gal4* (Jenett et al., 2012), which expresses in a subset of neurons projecting into the dorsal fan-shaped body (dFB), the sleep control center in *Drosophila* (Donlea et al., 2011; Pimentel et al., 2016). Using the GR23E10-*gal4* driver, we expressed human *dec2^P384R^* and *dec2^WT^* (as a control for *dec2* over-expression) in *Drosophila* dFB sleep neurons and assessed the effect on sleep. Hereafter, WT refers to GR23E10-*gal4*/+ control, while *dec2^WT^* and *dec2^P384R^* refer to the respective *dec2* transgenes over-expressed in GR23E10-*gal4* specific neurons. Although no significant differences were observed during nighttime sleep between the three groups, *dec2^P384R^* mutant flies displayed significantly shorter daytime sleep compared to WT and *dec2^WT^* (Fig.1A-1C). Expression of *dec2^P384R^* also reduced the average sleep bout duration while concomitantly increasing the sleep bout number, (Fig. 1D and 1E), suggesting that sleep is less consolidated in *dec2^P384R^* mutants.

**Figure 1:**
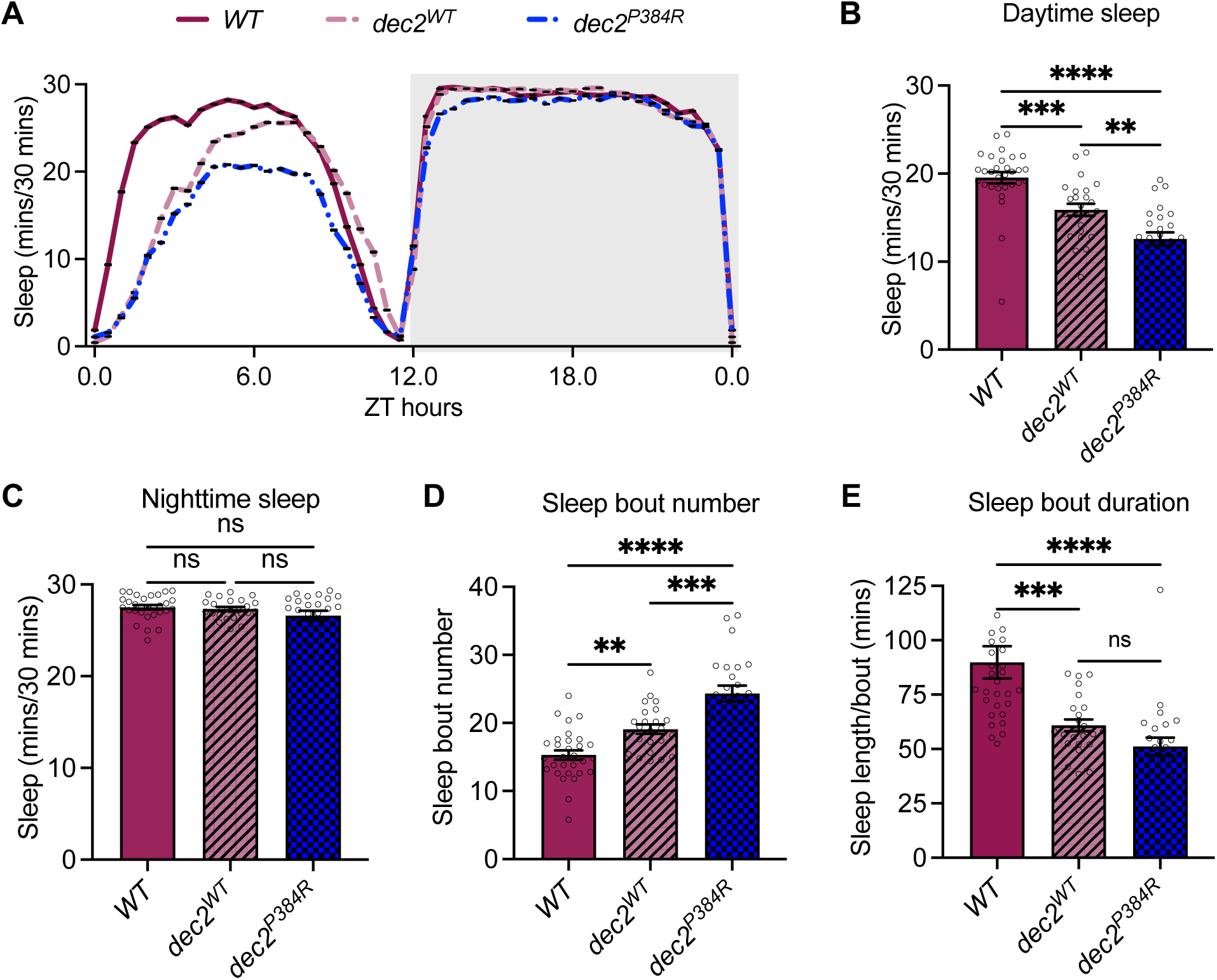
Expression of human *dec2P384R* in *Drosophila* sleep neurons reduces sleep. **A.** Sleep analysis in 12:12h L:D condition for *WT* (n=30), *dec2WT* (n=24), and *dec2P384R* (n=24) genotypes. **B-C.** Average sleep during daytime (B) and nighttime (C) of the genotypes indicated. **D-E.** Average sleep bout number (D) and sleep length/bout (E) in the genotypes indicated. ns=not significant, *p<0.05, **p<0.01, ***p<0.001, ****p<0.0001; one-way ANOVA with Tukey’s multiple comparisons.

Typically, aging is associated with deregulation of circadian rhythms and sleep homeostasis leading to fragmented sleep patterns, short nocturnal sleep duration, and reduced slow-wave sleep (Li et al., 2018). Because we observed fragmented sleep patterns in young *dec2^P384R^* mutants, we examined whether aging further impacted their sleep architecture. Consistent with previous studies, 60-day-old control flies showed more fragmented sleep compared to young flies (Fig. S1A-G). Specifically, sleep bout number increased by 148.5 % in old vs. young control flies (Fig. S1F). In *dec2^P384R^*mutants, sleep fragmentation also increased with age, albeit to a lesser degree (69.66 % increase in sleep bout number in old vs. young *dec2^P384R^* mutant flies). Nevertheless, these data indicate that *dec2^P384R^* mutants still exhibit age-dependent changes in sleep.

We also examined the effect of *dec2^P384R^*expression on sleep homeostasis, a regulatory mechanism that governs the timing and amount of sleep in a 24hr circadian period (Deboer, 2018). Normally, sleep pressure, or the drive to sleep, increases when animals are awake and decreases as animals sleep. Moreover, sleep deprivation further elevates sleep pressure and promotes longer periods of sleep in the next cycle to compensate for prior sleep loss (i.e., sleep rebound) (Deboer, 2013; Shaw et al., 2000). To determine the effect of *dec2^P384R^* expression on sleep homeostasis, we examined the total amount of sleep recovery after 24 hours of sleep deprivation. Control flies displayed a typical increase in sleep in the immediate cycle following the deprivation; however, *dec2^P384R^* mutants resumed a sleep pattern that was not significantly different from the pre-deprivation state (Fig. 2A and 2B). Additionally, control flies also displayed longer sleep bout duration with fewer sleep bout number indicating more consolidated sleep after sleep deprivation, whereas *dec2^P384R^* mutants did not display a significant change in sleep consolidation (Fig. 2C and 2D). Thus, the *dec2^P384R^* mutation interferes with natural sleep homeostasis. Collectively, we have established a *Drosophila dec2^P384R^*model that mirrors the mammalian short sleep phenotype.

**Figure 2:**
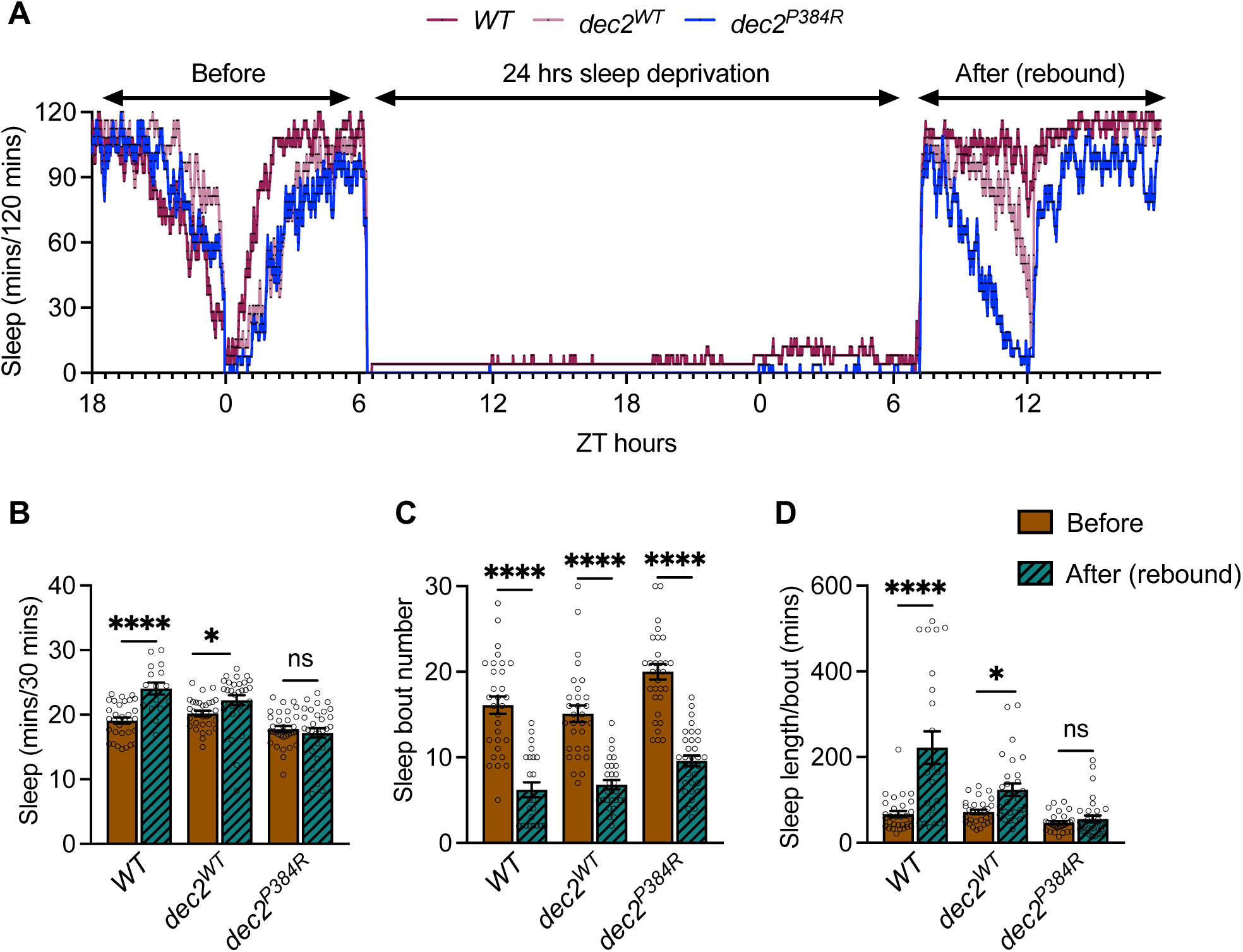
Expression of human *dec2P384R* in *Drosophila* sleep neurons reduces sleep rebound after sleep deprivation (SD). **A.** Sleep profiles for *WT, dec2WT* and *dec2P384R* genotypes before and after 24 hrs of SD. **B.** Average total sleep before and after SD for the genotypes indicated. **C.** Average sleep bout number before and after SD for the genotypes indicated. **D.** Average sleep length/bout before and after SD for the genotypes indicated. *WT* before (n=32) and after SD (n=16); *dec2WT* before (n=31) and after SD (n=30); *dec2P384R* before (n= 32) and after SD (n=32). ns=not significant, *p<0.01, ****p<0.0001; two-way ANOVA with Tukey’s multiple comparisons.

### dec2^P384R^ short sleep mutants live longer with improved health

In multiple animal models, complete loss of sleep decreases lifespan. For example, *Drosophila sleepless* mutants display 80% sleep loss and show a >50% reduction in lifespan (Koh et al., 2008). In humans, patients with a rare genetic disease called Fatal Familial Insomnia lose the ability to sleep around mid-life and only survive on average 18 months after diagnosis (Medori et al., 1992). In less severe instances, chronic sleep deprivation is associated with developmental disorders, cognitive impairments, metabolic dysfunctions, physiological deficits, cardiovascular diseases, and neurodegenerative diseases (Cappuccio et al., 2011; Deboer, 2013; Gambetti et al., 1995; Harper et al., 2001; Laaboub et al., 2022; Mazzotti et al., 2014; Mirmiran et al., 1983; Scullin and Bliwise, 2015; Sejbuk et al., 2022; Zhai et al., 2015). This prompted us to explore whether chronic reduced sleep in the *dec2^P384R^*mutants had any negative health impacts. Remarkably, we found that mutant *dec2^P384R^*flies lived significantly longer compared to control flies (Fig. 3A and Fig. S8). Thus, despite sleeping less, the *dec2^P384R^*mutation might in fact confer longevity to the organism.

**Figure 3:**
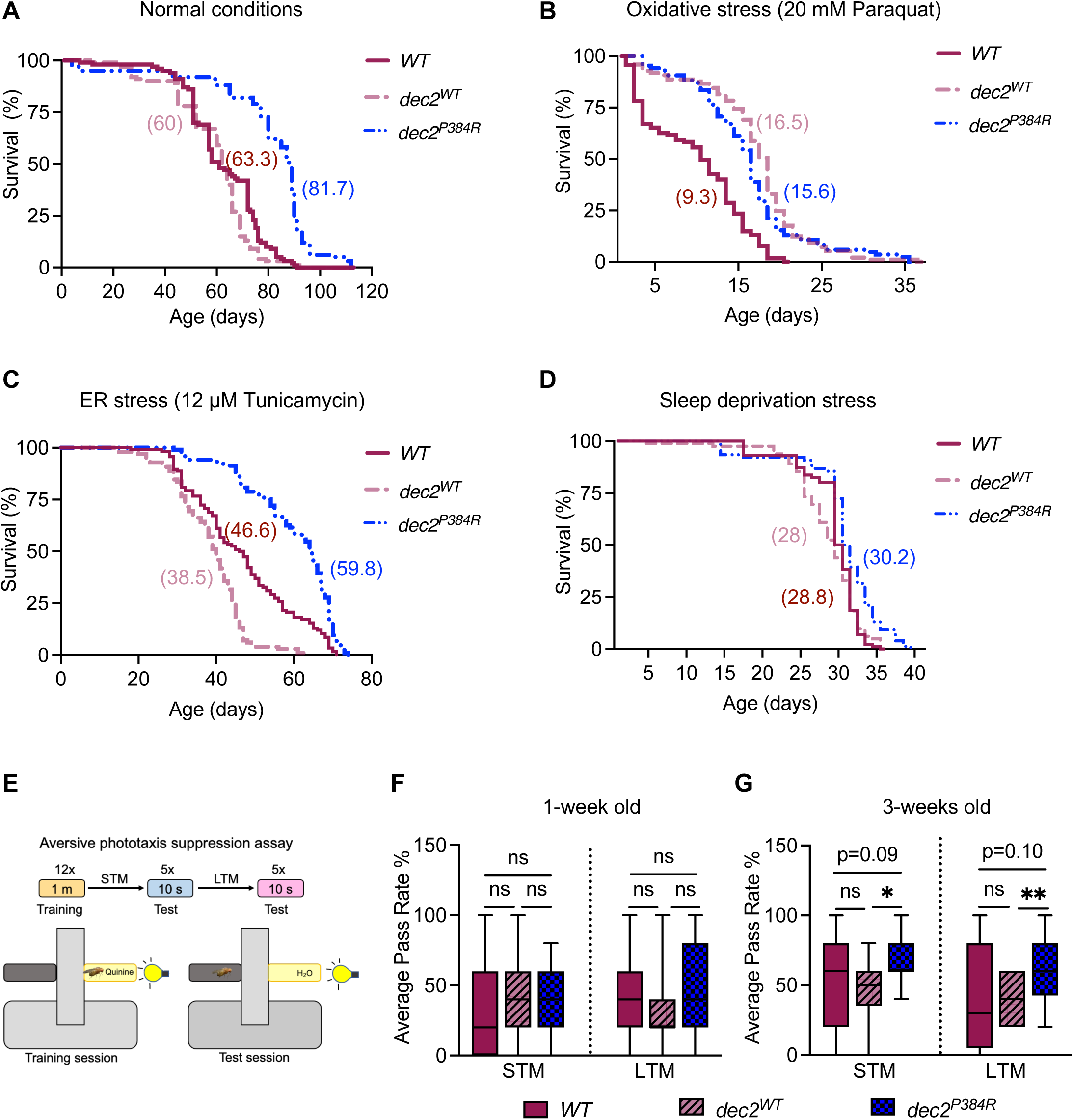
*dec2P384R* short sleep mutants live longer with improved health. **A-D.** Lifespan analysis of *WT, dec2WT and dec2P384R* genotypes under normal conditions (A), fed 20 mM Paraquat to induce oxidative stress (B), 12 μM Tunicamycin to induce ER stress (C) or under sleep deprivation stress (D). See Table S2 for descriptive statistics and log-rank test results. **E.** Schematic of aversive phototaxis suppression assay. **F-G.** Aversive phototaxis suppression assay of *WT, dec2WT and dec2P384R* genotypes for short term memory (STM) and long-term memory (LTM) at one week (F) and three weeks (G) of age (n=21). ns=not significant, *p<0.05, **p<0.01; two-way ANOVA with Tukey’s multiple comparisons.

To investigate whether *dec2^P384R^* mutants have improved physiology, we assessed health parameters that are often jeopardized with chronic sleep loss to determine if the *dec2^P384R^* mutation improves health. Sleep loss elevates the accumulation of reactive oxygen species (ROS) leading to oxidative stress in mice and flies, and if prolonged, reduces lifespan (Vaccaro et al., 2020). Similarly, sleep loss correlates with increased ER stress response pathways in mice (Mackiewicz et al., 2007; Naidoo et al., 2005; Terao et al., 2003) and *Drosophila* (Shaw et al., 2000), suggesting that sleep loss leads to ER stress. Therefore, we examined survival under these two stressors. First, flies were fed Paraquat, an organic cation that models oxidative stress via NADPH-dependent production of superoxide, ROS (Cochemé and Murphy, 2008). The lifespan of *dec2^P384R^* mutants was significantly longer than WT under oxidative stress conditions, indicating that *dec2^P384R^* mutants are resistant to oxidative stress (Fig. 3B and Fig. S8). Overexpression of *dec2^WT^* also improved survival under oxidative stress, suggesting that expression of the WT *dec2* gene confers some resistance to oxidative stress as well. *dec2^P384R^*mutants also displayed increased resistance against the ER stressor, Tunicamycin, which inhibits protein glycosylation in the ER, leading to accumulated unfolded proteins (Brown et al., 2014; Bull and Thiede, 2012) (Fig. 3C and Fig. S8). Additionally, we examined the sensitivity of *dec2^P384R^* mutant flies to sleep deprivation. Using the Sleep Nullifying Apparatus (SNAP) (Melnattur et al., 2020), flies were subjected to constant mechanical sleep deprivation and their lifespan was recorded. Consistent with increased stress resistance, *dec2^P384R^*mutants lived significantly longer than *WT* and *dec2^WT^* under sleep deprivation conditions (Fig. 3D and Fig. S8).

Sleep deprivation has also been linked to poor memory consolidation; in flies, 6-12 hours of sleep deprivation is sufficient to cause learning impairment (Seugnet et al., 2008). This suggests that altered sleep architecture can negatively impact memory encoding. Therefore, we examined whether *dec2^P384R^* mutants displayed significant memory impairment in either early-and/or mid-age by performing an aversive phototaxis suppression (APS) assay (Fig. 3E), which is commonly used to assess short and long-term memory in *Drosophila* (Ali et al., 2011; Seugnet et al., 2009). At one-week of age, there was no significant difference among mutants and controls (Fig. 3F). However, at three weeks of age (mid-life), *dec2^P384R^* mutants displayed significantly improved short and long-term memory compared to both control groups (Fig. 3G). Thus, mid-life memory function of *dec2^P384R^* mutants is in fact improved. Collectively, these data indicate that *dec2^P384R^* mutants live longer with improved health, despite sleeping less.

### dec2^P384R^ mutants exhibit improved mitochondrial capacity in flight muscles

Mitochondria are critical regulators of cellular energy and metabolism (Spinelli and Haigis, 2018), and improving mitochondrial function, either via increasing respiratory capacity or increasing biogenesis, can extend lifespan (Dillon et al., 2012; Ferguson et al., 2005; Nicholls, 2002; Nisoli et al., 2005; Ocampo et al., 2012). Moreover, clock rhythmicity determines energetic potential by signaling a need for reducing equivalents to drive oxidative phosphorylation (OXHPOS) (Aguilar-López et al., 2020; Masri et al., 2013). To determine if *dec2^P384R^* mutants display altered energy production, we first measured mitochondrial respiratory fluxes across the primary substrate-coupling pathways in homogenized flight muscles (Fig. 4A and 4B). *dec2^P384R^*mutants exhibited normal OXPHOS capacity supported by nicotinamide adenine dinucleotide (NADH) linked substrates, Proline and complex IV (Fig. 4C,4D and 4H). Strikingly, there was a significant increase in OXPHOS capacity supported by flavin adenine dinucleotide (FAD) linked substrates, namely succinate (61% vs. *dec2^WT^*, 81% vs. WT) and glycerol-3-phosphate (25% vs. *dec2^WT^*, 47% vs. WT) in *dec2^P384R^*relative to both WT and *dec2^WT^* flies (Fig. 4E-4G). Consistently, the FAD pool was also depleted in *dec2^P384R^* flies relative to both controls (Fig. 4I), indicative of decreased FAD/FADH_2_ ratio, favoring oxygen consumption and ATP synthesis. The increased respiratory flux observed in *dec2^P384R^* mutants was not attributed to a change in mitochondrial supercomplex structure and formation in *dec2^P384R^*and *dec2^WT^* flies (Fig. S2A); however, we found a significant decrease in citrate synthase activity, a marker of mitochondrial abundance, in *dec2^P384R^* flight muscles compared to controls (Fig. S2B). Thus, it is even more remarkable that *dec2^P384R^* mutants have improved mitochondrial capacity despite having less mitochondrial content. These data suggest that *dec2^P384R^* mutants exhibit improved mitochondrial respiratory function and ATP production capacity of substrates linked to reduction of FAD.

**Figure 4:**
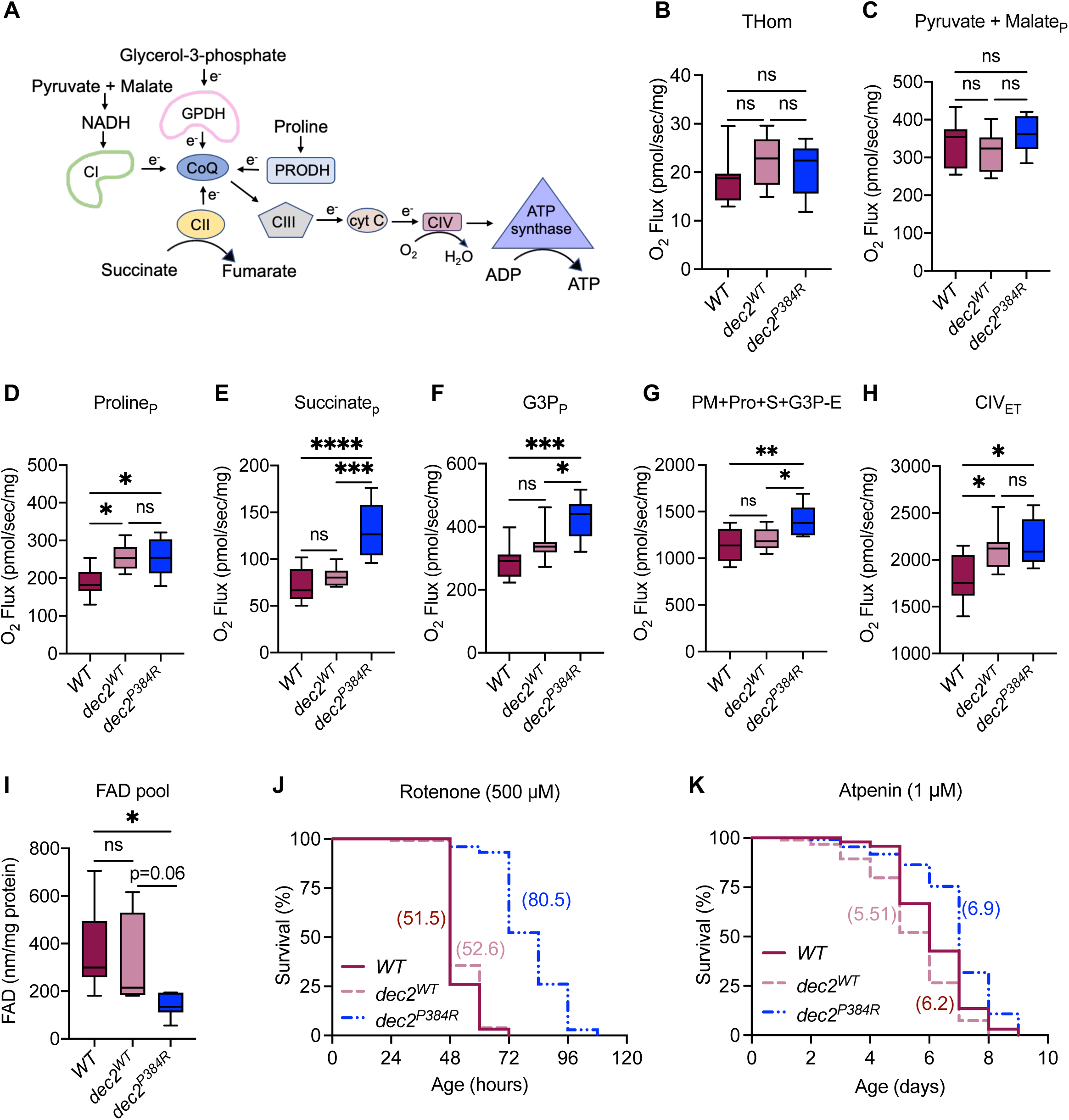
*dec2P384R* mutants exhibit improved mitochondrial capacity in flight muscles. **A.** Schematic illustration of substrate coupling to mitochondrial respiratory pathways evaluated by high-resolution respirometry. **B-G.** Respiration supported by the indicated substrates in the presence of ADP for *WT, dec2WT and dec2P384R* genotypes (THom, tissue homogenate). **H.** Respiration supported by complex IV in the presence of FCCP (ET, electron transfer). **I.** FAD pool. ns=not significant, *p<0.05, ***p<0.001, ****p<0.0001; one-way ANOVA with Tukey’s multiple comparisons. **J-K.** Lifespan of the genotypes indicated fed high doses of Rotenone (500 µM) (J) or Atpenin (1 µM) (K). See Table S2 for descriptive statistics and log-rank test results.

Based upon our observations that *dec2^P384R^* mutants display enhanced mitochondrial functional capacity, we tested resistance to stress induced by complex-specific OXPHOS inhibitors. We found that *dec2^P384R^* mutants survived significantly longer when fed high doses (500µM) of Rotenone (Fig. 4J and Fig. S8), a potent inhibitor of NADH oxidation and complex I activity. However, *dec2^P384R^* mutants fed high doses of the complex II inhibitor, Atpenin A5 (1µM) demonstrated a less pronounced improved survival (Fig. 4K and Fig. S8), consistent with *dec2^P384R^* mutants acting on FAD-linked substrate coupling. Taken together, these results indicate that *dec2^P384R^*mutants have increased FAD-linked mitochondrial respiratory capacity, which confers stress resistance and contributes to improved survival.

### Multiple stress response genes are upregulated in dec2^P384R^ mutants

Given that Dec2 is a transcription factor, we hypothesized that the increased lifespan and improved health of *dec2^P384R^* mutants might be due to global changes in gene expression. To examine this possibility, we performed Illumina-based RNA-sequencing to identify differentially expressed genes (DEGs) in *dec2^P384R^* vs. WT and *dec2^WT^* flies. One week old flies were collected at ZT3, the time in which we observed the most significant difference in their daytime sleep (Fig. 1A), and RNA was extracted from whole animals for sequencing. In parallel, we also performed long-read sequencing using Nanopore technology, which enables whole transcript sequencing and can identify isoform variants and limits amplification biases (Wang et al., 2021). Significantly, the two analyses shared ∼50% overlap in the DEGs identified (Fig. 5A), underscoring the confidence and reproducibility of our datasets. Principal component analyses were plotted to visualize the difference in gene expression among the three groups: *WT, dec2^WT^*and *dec2^P384R^* (Fig. S3A). The Illumina analyses obtained RNA-seq profiles for 17,972 genes with 323 DEGs in *dec2^P384R^* vs *WT* and 121 DEGs in *dec2^P384R^*vs *dec2^WT^* (Fig. 5B and 5C), while the long-read Nanopore sequencing obtained RNA-seq profiles for 15,488 genes with 136 DEGs in *dec2^P384R^*vs *WT* and 43 DEGs in *dec2^P384R^* vs *dec2^WT^*(Fig. S3B and S3C). To begin deciphering the molecular pathways that may be contributing to the improved health and extended lifespan of *dec2^P384R^*mutants, we performed gene ontology (GO) and KEGG pathway enrichment analyses (Fig. S4A and S4B). Notably, multiple gene clusters related to stress resistance were upregulated in *dec2^P384R^* mutants (Fig. 5D), which could account for the improved physiological health observed in *dec2^P384R^*mutants. Moreover, we identified several uncharacterized and orphan genes that were differentially expressed in *dec2^P384R^* mutants (Fig. 5B-5E), which could represent novel pro-longevity factors.

**Figure 5:**
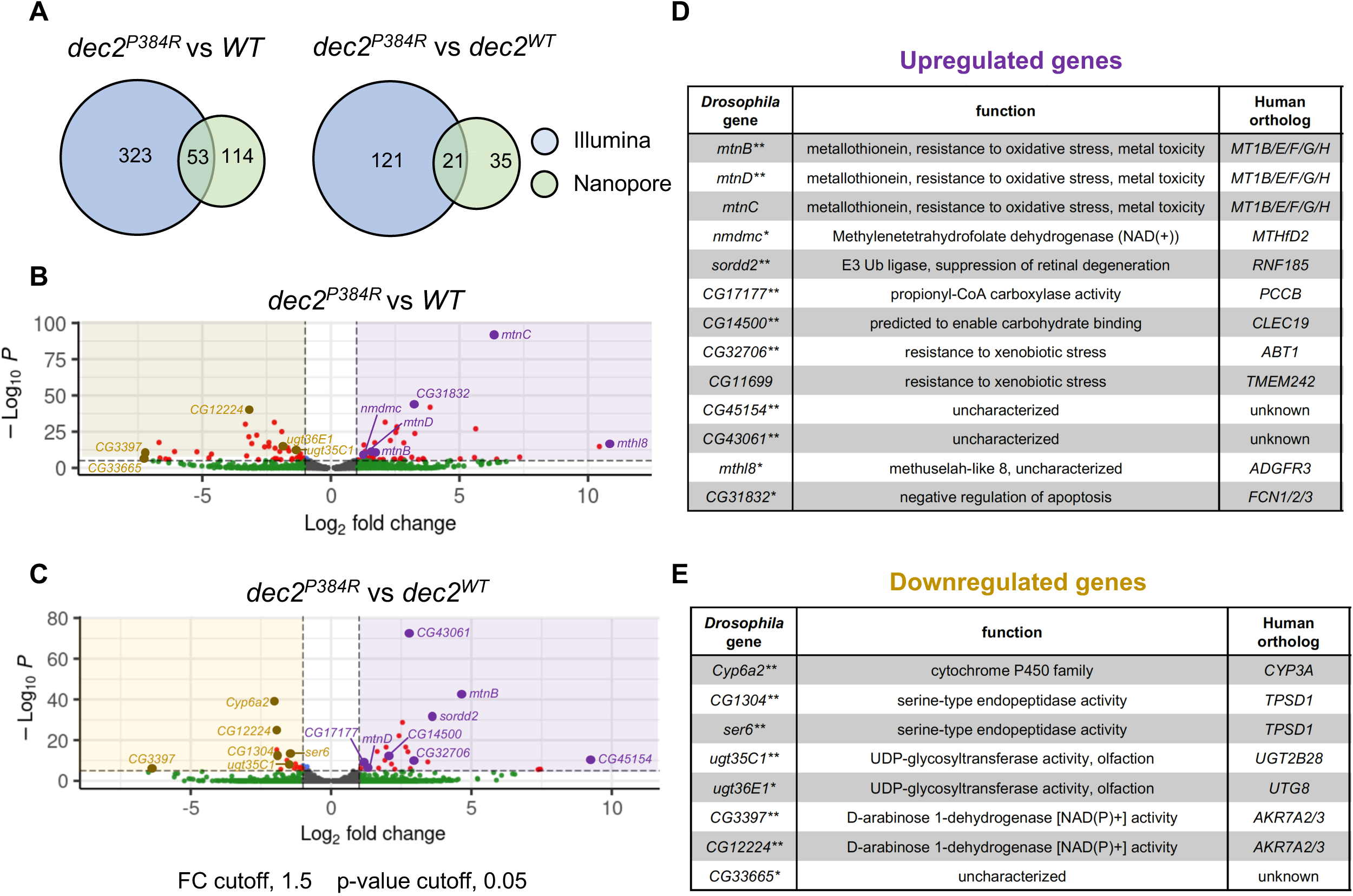
Multiple stress response genes are upregulated in *dec2P384R* mutants. **A**. Venn diagrams comparing DEGs identified in Illumina and Nanopore analyses. **B-C**. Volcano plots of DEGs in *dec2P384R* vs. *WT* (B) and *dec2P384R* vs. *dec2WT* (C) identified in the Illumina analyses. Significantly down-regulated genes are on negative side (left), significantly up-regulated genes are on positive side (right). Cutoff ranges: log fold changes of −1.5 and +1.5; padj-value of 0.05. **D-E.** Table of upregulated (D) and downregulated (E) genes of interest, *identified in both Illumina and Nanopore analyses as a DEG in *dec2P384R* vs. *WT*, **identified in both Illumina and Nanopore analyses as a DEG in *dec2P384R* vs. *dec2WT.* See Tables S3 and S4 for full gene expression profiles obtain from the Illumina (Table S3) and Nanopore (Table S4) analyses.

To validate the RNA-seq data, we selected the top ten upregulated genes as well as a subset of related gene family members to quantify expression by qPCR (Fig. 6A-6C and S5A-S5F). We first examined the metallothionein (MT) gene family, which consists of five paralogs (*mtnA-E*) that reside in a gene cluster on Ch. 3R (Fig. 6A). MT proteins have known cytoprotective functions and promote cell survival with increased expression (Bakka et al., 1982; Dutsch-Wicherek et al., 2008; Iszard et al., 1995; Sato and Bremner, 1993). Consistent with the RNA-seq data, *mtnB* and *mtnD* transcripts were increased in *dec2^P384R^* compared to both controls (Fig. 6A). *Methuselah-like 8* (*mthl8*) is an uncharacterized gene that is predicted to encode a G protein-coupled receptor (Harmar, 2001) and was the most upregulated gene in *dec2^P384R^*vs *WT* in both Illumina and Nanopore datasets (Fig. 5B and S3B). Notably, a related homolog *methuselah* has been linked to lifespan regulation in flies (Lin et al., 1998). Strikingly, *mthl8* transcripts measured by qPCR were increased >1000-fold in both *dec2^P384R^* and *dec2^WT^*compared to *WT* (Fig. 6B). Finally, we examined expression of *CG11699*, which was upregulated in *dec2^P384R^* compared to both controls; *CG11699* transcripts were increased in *dec2^WT^* compared to *WT* and further upregulated in *dec2^P384R^* (Fig. 6C). Although *CG11699* is not well-characterized, it has been linked to lifespan regulation; a transposable element insertion in the 3’UTR increases *CG11699* expression and extends lifespan (Mateo et al., 2014). *CG11699* is also related to human TMEM242, a component of the mitochondrial proton-transporting ATP synthase complex (Carroll et al., 2021). Overall, these results are consistent with the RNA-seq data and hint that multiple stress response pathways are upregulated in *dec2^P384R^* mutants.

**Figure 6:**
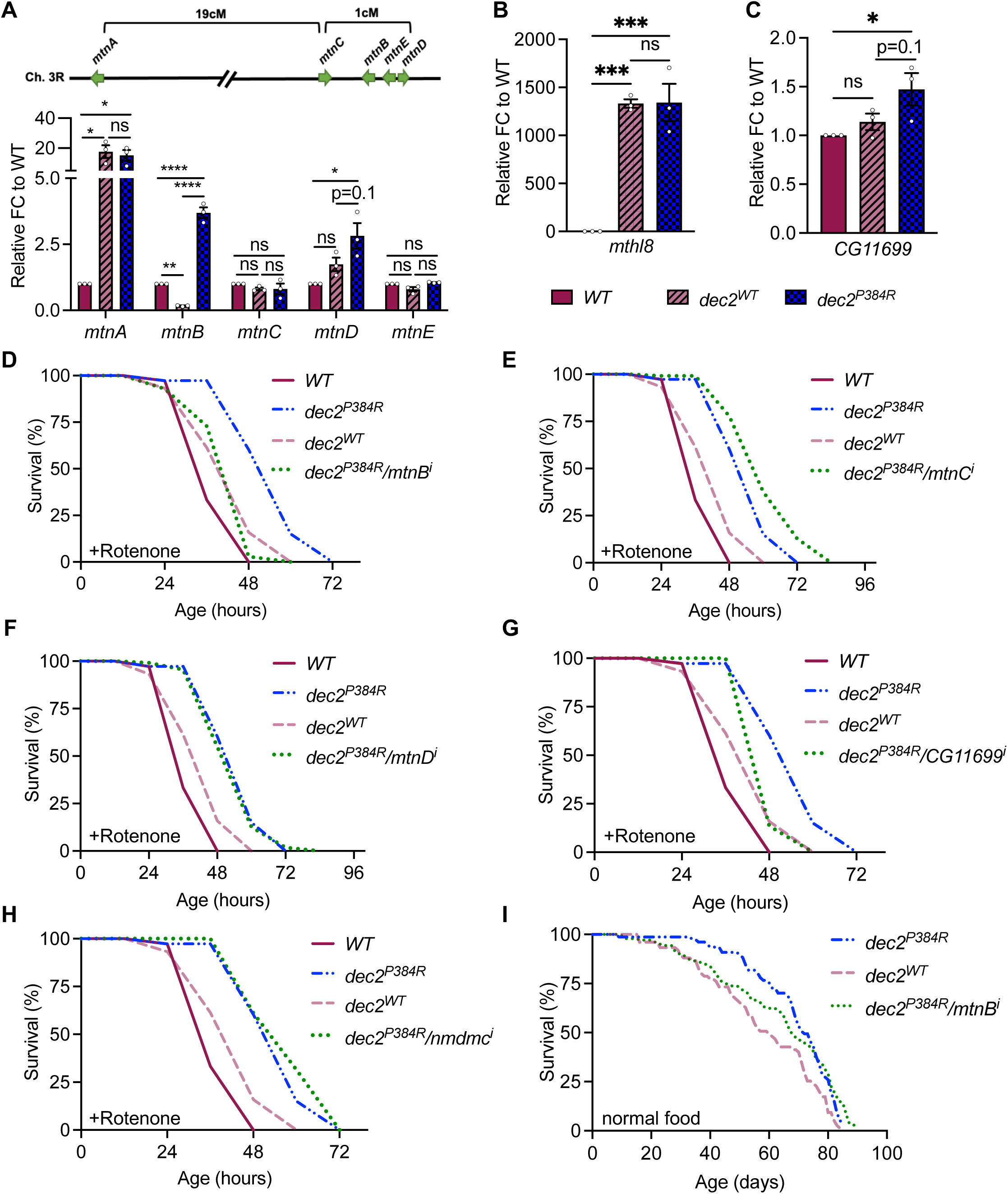
Lifespan extension in *dec2P384R* mutants is dependent on increased *mtnB* expression. **A.** Schematic of *mtn* gene cluster on Ch. 3R (top) and expression of *mtn* genes measured by qPCR (bottom). **B-C.** Expression of *mthl8* (B) and *CG11699* (C) measured by qPCR. FC=fold change, (n=3), ns=not significant, *p<0.05, **p<0.01, ***p<0.001, ****p<0.0001; one-way ANOVA with Tukey’s comparisons. **D-H.** Lifespan of flies fed 500 µM Rotenone. **I.** Lifespan of flies under normal conditions. See Table S2 for full descriptive statistics and results of log-rank tests.

### Lifespan extension in dec2^P384R^ mutants is dependent on increased mtnB expression

To identify which gene(s) are critical for regulating the lifespan extension of *dec2^P384R^* mutants, we examined whether inhibition of any of the top upregulated genes identified in the differential gene expression (DGE) analyses could negate the lifespan extension of *dec2^P384R^* mutants. To do this, we utilized two complementary binary expression systems, LexAop/LexA and UAS/GAL4, to simultaneously express *dec2^P384R^* in sleep neurons and inhibit candidate gene expression via RNAi in various tissues, respectively. We first tested whether the GR23E10-*lexA*/*lexAop-dec2^P384R^* transgenic expression system induced a short-sleep phenotype like the GAL4/UAS system and, indeed, we observed a similar short sleep phenotype when *dec2^P384R^* was expressed using the LexA/LexAop system (Fig. S6A-S6E).

GR23E10-*lexA*/*lexAop-dec2^P384R^* transgenic flies were also resistant to Rotenone (Fig. 6D and Fig. S8) but still do not survive for more than three days. Therefore, we performed lifespans in the presence of Rotenone as a faster means of screening through candidate genes initially. We first examined the *MT* gene family *mtnA-E*. Upregulation of *MT* genes in neurons promotes longevity (Bahadorani et al., 2010); thus, we used the pan-neuronal driver *elav-gal4 to* inhibit *MT* gene expression in the brain of *dec2^P384R^* flies. Strikingly, inhibition of *mtnB* alone was sufficient to diminish the lifespan extension effect of *dec2^P384R^*mutants back to control lifespans (Fig. 6D and Fig. S8), while there was no significant difference in lifespan with suppression of *mtnC* or *mtnD* (Fig. 6E, 6F and Fig. S8). This is consistent with the DGE analyses, as *mtnB* was the most differentially expressed MT gene compared to both WT and *dec2^WT^* controls (Fig. 5B and 5C). We next tested *CG11699* and *nmdmc* by inhibiting their expression ubiquitously using tubulin-*gal4* (*tub-gal4*). However, we did not obtain any viable progeny, suggesting that global inhibition of these genes is lethal. We then inhibited *CG11699* or *nmdmc* in neurons using *elav*-*gal4*. Inhibiting *CG11699* in *dec2^P384R^* mutants reduced the lifespan back to WT (Fig. 6G and Fig. S8), while there was no reduction in lifespan with *nmdmc* gene suppression (Fig. 6H and Fig. S8). Finally, we examined whether *mtnB* is required for *dec2^P384R^* mutant lifespan extension under normal conditions and found that inhibition of *mtnB* also partially reduced the lifespan of *dec2^P384R^* mutants under normal conditions (Fig. 6I and Fig. S8); thus, *mtnB* expression in neurons significantly contributes to the lifespan extending effects of *dec2^P384R^* expression. Collectively, we have identified at least two critical factors that are required for the lifespan extending effects of *dec2^P384R^*mutants and these data reinforce that the lifespan extension observed in *dec2^P384R^* is a result of increased expression of stress-response signaling pathways.

### Improved health correlates with reduced sleep

Although the *dec2^P384R^* mutation is known to promote prolonged wakefulness by increasing *orexin* expression in mammals (Hirano et al., 2018), it is puzzling that over-expression of mammalian *dec2^P384R^* in *Drosophila* can still induce a short-sleep phenotype given that the *orexin* system does not exist in invertebrates (Soya and Sakurai, 2020). This suggests that *dec2^P384R^* is capable of reducing sleep by an orexin-independent mechanism. This led us to postulate that perhaps the *dec2^P384R^*-dependent short sleep phenotype is not directly related to altered core sleep mechanisms, but rather a byproduct of their increased longevity. Based on this idea, we hypothesized that inhibiting the pro-health pathways triggered by Dec2^P384R^ would reverse the short sleep phenotype. Thus, we inhibited *mtnB* pan-neuronally in *dec2^P384R^*mutants, which reduces the lifespan of *dec2^P384R^* mutants back to WT (Fig. 6I), and assessed their sleep length. In accord with our hypothesis, we found that *mtnB* inhibition increased sleep of *dec2^P384R^* mutants back to WT levels (Fig. 7A-7B). Moreover, inhibition of *mtnB* also suppressed the sleep fragmentation phenotype of *dec2^P384R^* mutants (Fig. 7C-7D). Thus, these data suggest that the improved health of *dec2^P384R^* mutants may also contribute to the short sleep phenotype.

**Figure 7:**
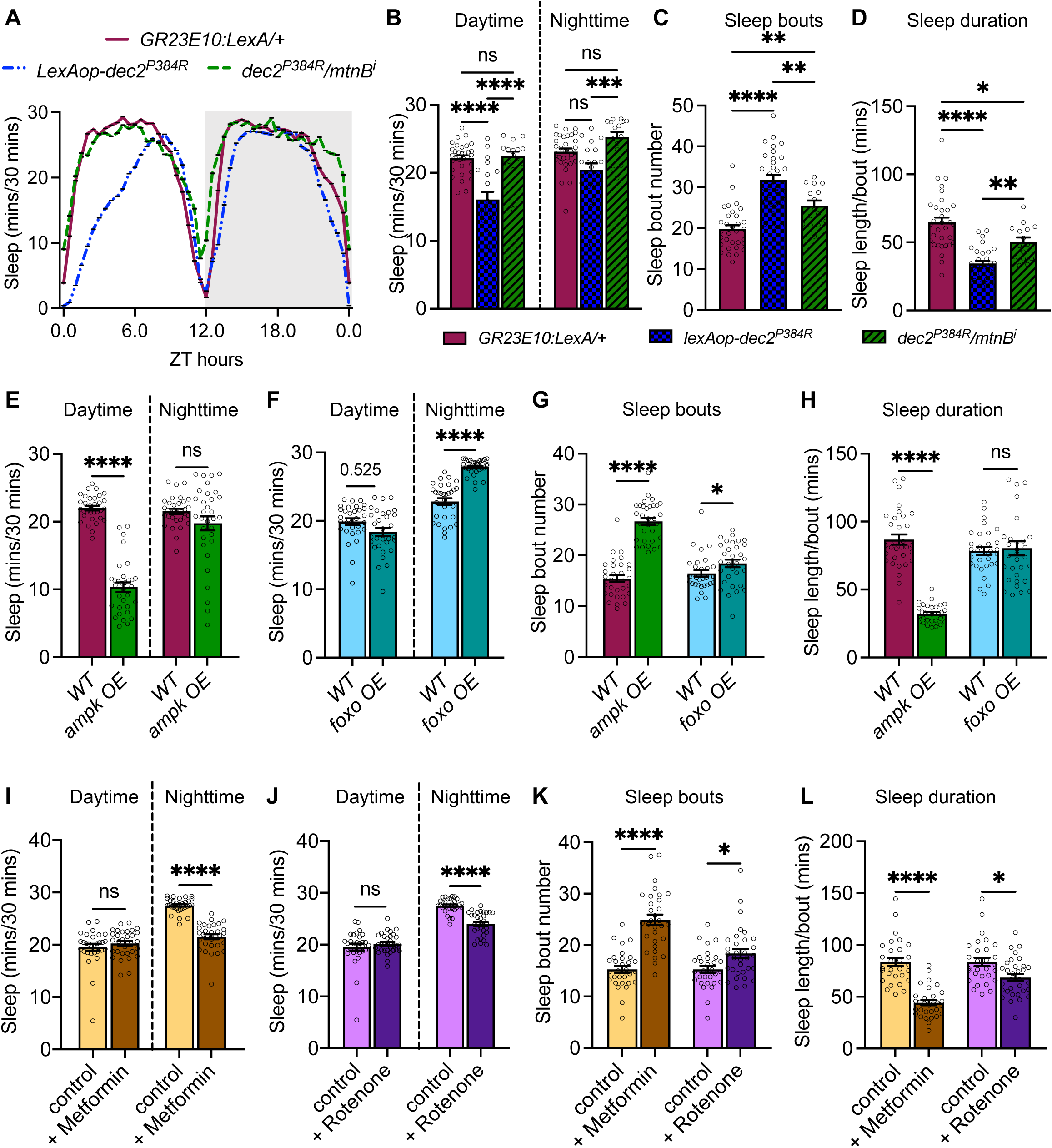
Improved health correlates with reduced sleep. **A.** Sleep analysis in 12:12h L:D condition for *GR23E10:LexA/+* (n=32), *lexAop-dec2P384R* (n=20) *and dec2P384R /mtnBi* genotypes (n=15). **B-D.** Average daytime sleep and nighttime sleep (B), sleep bout number (C) and sleep length/bout (D) for the genotypes indicated (ns=not significant, *p<0.05, ***p<0.001, ****p<0.0001). **E-F.** Average daytime sleep and nighttime sleep for long-lived mutants *ampk* OE (n=32) (E) and *foxo* OE (n=31) (F). **G-H.** Average sleep bout number (G) and sleep length/bout (H) for the long-lived mutants indicated. **I-J.** Average daytime and nighttime sleep for the long-lived models, 500 mM Metformin (n=31) (I) and 0.1 µM Rotenone (n=32) (J). **K-L.** Average sleep bout number (K) and sleep length/bout (L) for the long-lived models indicated (n=32), ns=not significant, **p<0.01, ***p<0.001, ****p<0.0001; one-way ANOVA with Šidák multiple comparisons.

Based on our result that reducing pro-health pathways in *dec2^P384R^* mutants reverses the short sleep phenotype, it is intriguing to speculate that improving organismal health may reduce sleep pressure. Although multiple studies have shown that aging correlates with increased sleep disturbances (De Nobrega and Lyons, 2020; Mattis and Sehgal, 2016), how activation of pro-longevity pathways affects sleep has not been explored as extensively. However, in a study using a sleep inbred panel, in which flies were sorted based on their natural sleep time, short sleep flies displayed 16% longer lifespan compared to long sleep flies (Thompson et al., 2020). Moreover, it has been demonstrated previously that reducing insulin signaling, which promotes longer lifespan (Hwangbo et al., 2004), also reduces daytime sleep (Metaxakis et al., 2014). Thus, there is compounding evidence to suggest that enhancing organismal health can reduce sleep pressure. We further explored this idea by assessing sleep in two other long-lived models: over-expression of *AMPK* pan-neuronally (Ulgherait et al., 2014) and *foxo* in the fat body (Hwangbo et al., 2004), both of which promote longer lifespan by inducing cell nonautonomous mechanisms. Consistent with our hypothesis, both long-lived models also exhibited reduced sleep (Fig. 7E and 7F). We also observed increased sleep fragmentation in both mutant models compared to controls (Fig. 7G and 7H). Interestingly, fat body overexpression of *foxo* displayed increased nighttime sleep, which is consistent with the previous observation that inhibiting insulin signaling reduces daytime sleep, but promotes a compensatory increase in nighttime sleep (Metaxakis et al., 2014). However, we did not observe similar compensatory increases in nighttime sleep of *AMPK* or *dec2^P384R^* models, suggesting that sleep may be differentially influenced in these models. Nevertheless, these results lend further support to the notion that inducing pro-longevity pathways may reduce sleep pressure.

Finally, we tested whether non-genetic means of promoting health could also induce changes to sleep in WT animals. Administering low doses of mitochondria-targeted agents, such as Rotenone and Metformin, can improve health and extend lifespan by eliciting hormetic responses (De Haes et al., 2014; Martin-Montalvo et al., 2013; Na et al., 2015; Na et al., 2013; Onken et al., 2022; Schmeisser et al., 2013; Yuyun et al., 2013). Therefore, we examined sleep in WT flies that were fed 0.1 µM of Rotenone or 5 mM of Metformin, which are the optimal doses required for improved health span (Na et al., 2015; Schmeisser et al., 2013). Consistent with our model that improved health reduces sleep pressure, we observed reduced nighttime sleep and increased sleep fragmentation in WT flies fed either low doses of Metformin or Rotenone (Fig. 7I-7L). Taken together, these data lend support to the idea that improving health might reduce sleep need.

## Discussion

In this study we identified a familial natural short sleep mutation as a pro-longevity factor. While it has been suggested that human natural short sleepers are able to thrive with chronic short sleep, this has never been directly tested experimentally. Using a *Drosophila* model, we have demonstrated for the first time that expression of the short sleep mutation *dec2^P384R^* in fact extends lifespan and promotes healthy aging. Moreover, we identified metabolic adaptations and genetic pathways under the influence of neuronal Dec2 that contribute to the increased lifespan and stress resistance observed in *dec2^P384R^* mutants. Namely, multiple pathways related to metabolic and xenobiotic stress response pathways were upregulated. Recently, other familial natural short sleep (FNSS) mutations have been discovered in the human ADRB1, NPSR1, and GRM1 genes (Shi et al., 2021; Xing et al., 2019). Whether the paradigms we have established for the Dec2 mutation extend to these other FNSS mutations remains to be determined, but these studies provide a foundation for further investigation into potential links between natural short sleep mutations and health span.

Although there are likely multiple genes that collectively contribute to the lifespan extension of *dec2^P384R^* mutants, we found that increased expression of *mtnB*, a metallothionein protein, is one critical gene required for the full lifespan extension effects of *dec2^P384R^*mutants. Metallothionein proteins are small proteins that mediates cellular stress responses and are linked to longevity (Ebadi et al., 2005; Yang et al., 2006). Notably, increased expression of metallothionein results in resistance to mitochondrial induced stress and prevention of apoptotic signaling (Bahadorani et al., 2010; Ebadi et al., 2005; Hands et al., 2010; Yang et al., 2006). This is consistent with our observations that *dec2^P384R^* mutants are resistant to mitochondrial inhibitors (Fig. 4J and 4K). We also observed upregulation of the *CG11699* gene, which transcribes a protein that is not fully characterized. However, in flies, increased expression of *CG11699* confers xenobiotic stress resistance through increased aldehyde dehydrogenase type III (ALDH-III) activity (Mateo et al., 2014). ALDH oxidizes aldehydes to non-toxic carboxylic acids mitigating both intrinsic and pathological cellular stress, thus promoting overall survival (Shortall et al., 2021). Additionally, a closely related human homolog of *CG11699*, TMEM242, is required for the assembly of the c-8 ring of human ATP synthase, which is essential for ATP production (Carroll et al., 2021). Consistently, we found that *dec2^P384R^* mutants have increased mitochondrial respiratory capacity (Fig. 4A-4I). Specifically, we found improved FAD-linked capacity with a concomitant decrease in the FAD-pool, indicating an overall increase in FAD oxidation and ATP production. While increased FAD oxidation can also result in increased oxidative stress, we have found that the *dec2^P384R^* mutants are able to capitalize on the increased capacity while mitigating the potential deleterious effects of oxidative stress through upregulation of multiple stress-response mechanisms.

Our results also indicate that expressing the *dec2^P384R^* mutation in neurons alters cellular physiology in other non-neuronal tissues, such as muscles (Fig. 4). These data suggest that Dec2^P384R^ triggers cellular responses in a cell non-autonomous manner to elicit systemic changes. How might this occur? Dec2 is a transcription factor that regulates multiple circadian genes (Sato et al., 2016), many of which are known to affect organismal health and survival. For example, inhibiting the *C. elegans period* ortholog *lin-42* suppresses autophagy and accelerates aging (Kalfalah et al., 2016). Likewise, null mutations in the *Drosophila period* ortholog reduce resistance to oxidative stress (Krishnan et al., 2009), while neuronal overexpression of *period* extends lifespan and confers stress resistance (Solovev et al., 2019). Thus, upregulation of *period* improves health and extends lifespan. In mammals, *per1* expression is activated by CLOCK/BMAL, which bind to an upstream E-box binding site to induce *per1* transcription (Gekakis et al., 1998). WT DEC2 competes with CLOCK/BMAL at the E-box binding site, leading to repressed *per1* expression (Honma et al., 2002); however, mutant DEC2^P384R^ has reduced affinity to E-box promoter sequences (Hirano et al., 2018). Thus, it is conceivable that Dec2^P384R^ could lead to increased *period* expression in sleep neurons, which could subsequently trigger downstream cell non-autonomous physiological changes. Although we did not observe any significant changes to *period* transcripts in our RNA-seq data, the gene expression changes could be isolated to sleep neurons, which may have been precluded by our whole animal analysis. Examining transcriptional changes specifically in sleep neurons will be important future steps to identify direct targets of mutant Dec2^P384R^.

The *Drosophila* genome encodes a single gene, *clockwork orange* (*cwo*), that is orthologous to mammalian *dec1* and *dec2* (Kadener et al., 2007). Similar to DEC proteins, CWO also antagonizes CLOCK/BMAL transcription factors at an E-box site to attenuate *period* expression (Zhou et al., 2016). Although CWO is structurally similar to Dec2, containing a basic helix-loop helix domain, there is less than 18% amino acid sequence similarity with Dec proteins, and the proline 384 residue is not conserved in the CWO protein. Thus, there are likely to be functional distinctions between the orthologs. Nevertheless, the fact that mammalian *dec2^P384R^*induces short sleep and impacts multiple aspects of physiology when expressed in flies, signifies that it is acting in a dominant negative fashion and could interfere with expression of endogenous CWO target genes. Alternatively, the proline mutation could produce a more dramatic structural alteration to Dec2, causing it to bind ectopic sites in the genome and alter transcription of non-native CWO target genes. Having a deeper understanding of endogenous Dec2 and CWO target genes, perhaps with a focus on non-circadian regulatory networks, will be important to decipher how *dec2* orthologs and their variants influence non-sleep phenotypes.

Finally, our results also suggest that the improved health in *dec2^P384R^*mutants may also contribute to the short sleep phenotype. Typically sleep loss is associated with reduced health and lifespan (Gonzalez-Ortiz et al., 2000; Koh et al., 2008; Spiegel et al., 1999); however, there is evidence to suggest that this may not always be the case. In a study using a sleep inbred panel in which flies were sorted based on their natural sleep time, short sleep flies lived significantly longer compared to flies that slept longer (Thompson et al., 2020). This suggests that shorter sleep does not always strictly correlate with reduced lifespan. Moreover, this study and other previous studies (Metaxakis et al., 2014) have demonstrated that multiple long-lived mutants also sleep less, which leads to an intriguing question: does promoting longevity reduce sleep need? The fact that some species evolved mechanisms to virtually eliminate the need for sleep, while maintaining a similar lifespan as related species that require sleep (Duboué et al., 2011), lends support to this idea. Perhaps these species have naturally adapted sleep-independent pro-health mechanisms that allows them to survive with less sleep. This might also help explain why expression of the mammalian *dec2^P384R^* transgene can still induce a short sleep phenotype in flies, despite lacking an *orexin* ortholog. Thus, we hypothesize that the pro-health pathways that are ectopically induced in *dec2^P384R^*mutants may also contribute to the short sleep phenotype in flies. Whether similar mechanisms occur in mammals will be important future studies.

Sleep loss is becoming endemic in our modern society; it is estimated that 30% of adults in the U.S. sleep an average of 6hrs/night or less and are chronically sleep deprived (Schoenborn and Adams, 2010; Sheehan et al., 2019). These sleep disturbances are becoming even more prevalent due to certain occupational, and lifestyle demands (i.e., shiftwork, cross time-zone travel). Thus, sleep loss has become a major public health concern and uncovering mechanisms that can sustain health in sleep-deprived states is of critical importance. Studying the genetic mechanisms regulated by these rare short sleep mutations could provide a unique opportunity to not only understand how these exceptional individuals offset the negative effects of sleep deprivation, but also uncover novel pro-longevity pathways that could be co-opted to sustain health in sleep-deprived states as well as promote health more generally.

## Acknowledgements

We thank all lab members for critically reading the manuscript and providing helpful feedback.

## Author contributions

Conceptualization: A.E.J., S.R.L., P.P.; Methodology: P.P, S.R.L., E.R.M.Z., O.S.D., P.K.W.; Formal analysis: P.P., S.R.L., E.R.M.Z., O.S.D., P.K.W., C.L.A.; Investigation: P.P., S.R.L., E.R.M.Z., W.S.D. O.S.D., P.K.W., C.L.A., A.E.J.; Resources: O.S.D., P.K.W., J.P.K.; Data curation: P.P., S.R.L., E.R.M.Z., W.S.D. O.S.D., P.K.W., C.L.A., A.E.J.; Writing - original draft: A.E.J., P.P., P.K.W., E.R.M.Z., C.L.A.; Writing - review & editing: P.P., S.R.L., E.R.M.Z., W.S.D., J.P.K., O.S.D., P.K.W., C.L.A., A.E.J.; Supervision: A.E.J., C.L.A.; Project administration: A.E.J.; Funding acquisition: A.E.J., J.P.K., E.R.M.Z., S.R.L.

## Data Availability

All data are available in the main text or the supplementary materials. Additional information on data sources is available upon request from the corresponding author. All unique materials used in the study are available from the authors or from commercially available sources. For the gene expression analyses, the raw and processed data have been submitted to NCBI under the accession PRJNA957078. Data analysis code is available at github at https://github.com/pkerrwall/dec2_fly

