## Supplemental Figure for "A familial natural short sleep mutation promotes healthy aging and extends lifespan in *Drosophila*"

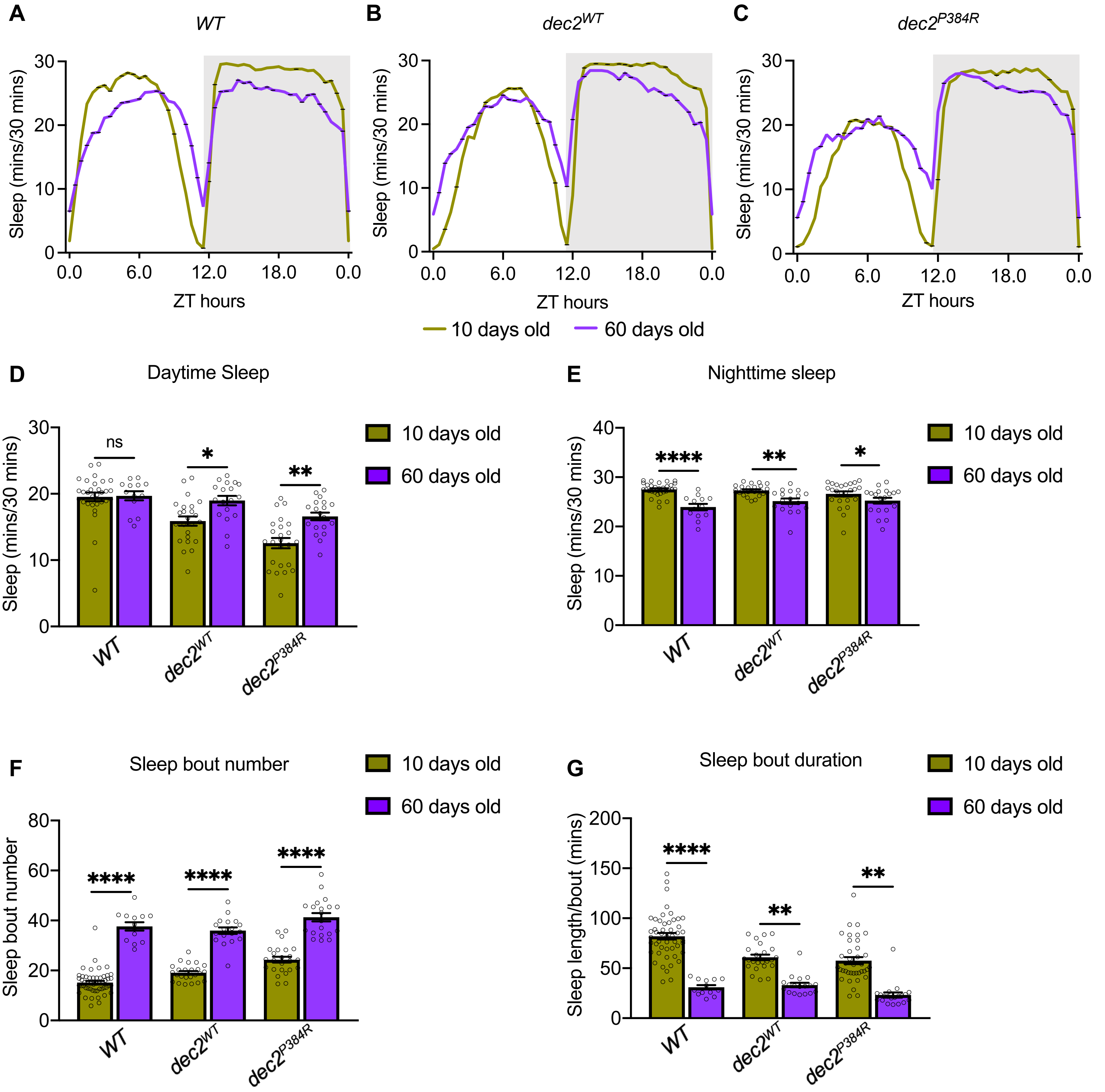


**Supplemental Figure 1: *dec2$^{P384R}$* mutants exhibit age-dependent changes in sleep. A-C.** Sleep analysis in 12:12h L:D condition at 10 and 60 days old for *WT* (A), *dec2$^{WT}$* (B) and *dec2$^{P384R}$* (C) genotypes. **D-E.** Average daytime (D) and nighttime (E) sleep for the genotypes indicated at 10 days and 60 days. ns=not significant, *p<0.05, **p<0.01, ****p<0.0001; one-way ANOVA with Tukey's comparisons. **F-G.** Average sleep bout number (F) and sleep length/bout (G) of the genotypes indicated at 10 days and 60 days. ns=not significant, **p<0.01, ****p<0.0001; one-way ANOVA with Tukey's comparisons. *WT* 10 days (n=30) and 60 days (n=13); *dec2$^{WT}$* 10 days (n=24) and 60 days (n=16); *dec2$^{P384R}$* 10 days (n=24) and 60 days (n=20).

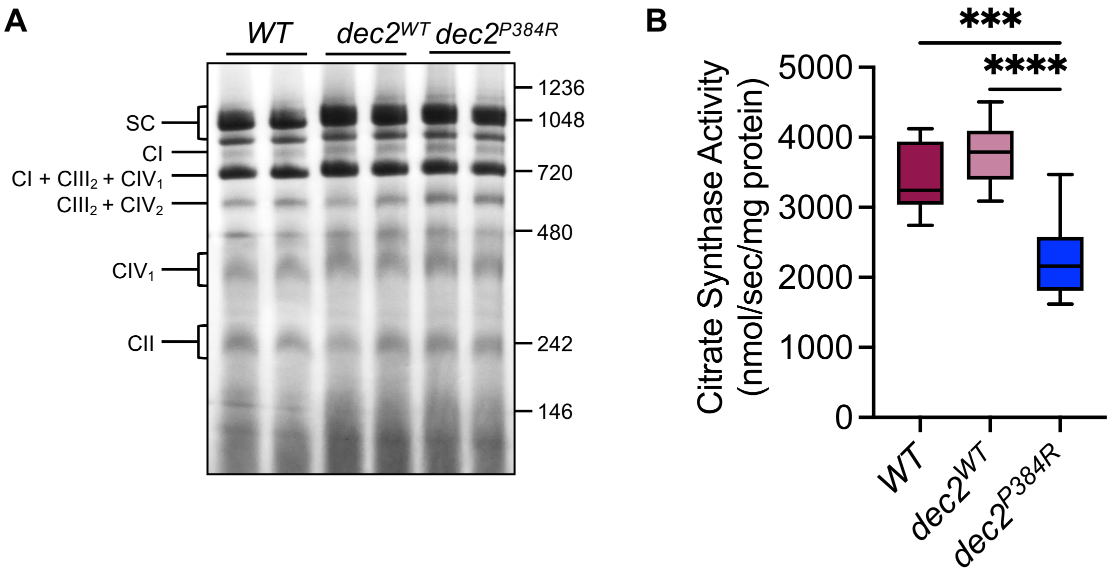


**Supplemental Figure 2: Supercomplex formation and citrate synthase activity. A.** Representative blue native gel of higher-order organization of the mitochondrial electron transport chain. **B.** Citrate synthase activity as a measure of relative mitochondrial abundance. ns=not significant, ***p<0.001, ****p<0.0001; one-way ANOVA with Tukey's multiple comparisons.

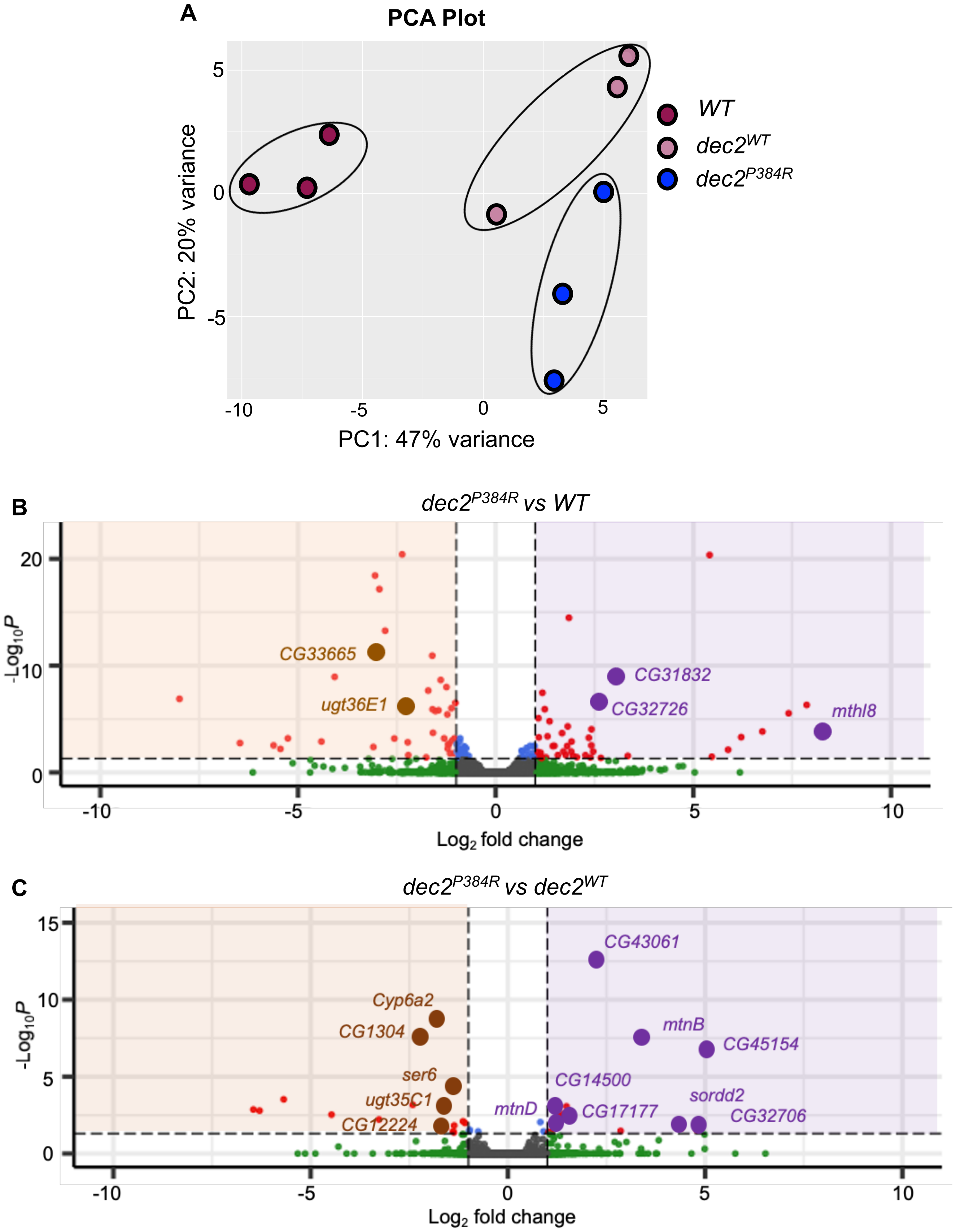


**Supplemental Figure 3: Differential gene expression analyses. A.** Principal Component Analysis (PCA) plot of Illumina-based gene expression among *WT, $dec2^{WT}$ and $dec2^{P384R}$* genotypes (n=3). **B-C**. Volcano plots of DEGs in *$dec2^{P384R}$* vs. *WT* (B) and *$dec2^{P384R}$* vs. *$dec2^{WT}$* (C) identified in the Nanopore analysis. Significantly down-regulated genes are on negative side (left), significantly up-regulated genes are on positive side (right). Cutoff ranges: log fold changes of -1.5 and +1.5 padj-value of 0.05.

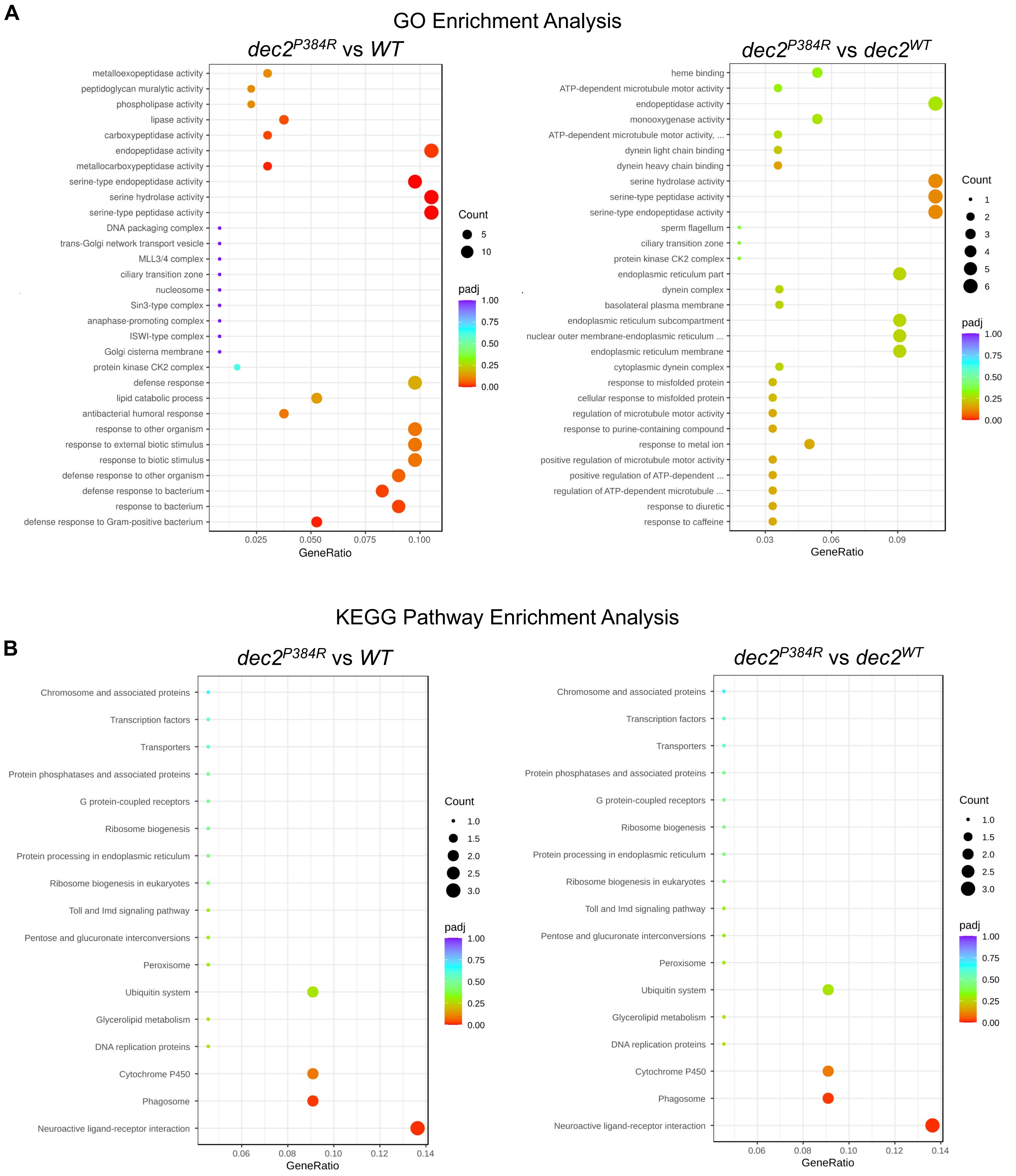


**Supplemental Figure 4: Enrichment analyses for the Illumina sequencing data. A-B.** Gene ontology (GO) enrichment scatter plot (A) and Kyoto Encyclopedia of Genes and Genomes (KEGG) pathway enrichment scatter plot (B) for *dec2*$^{P384R}$ vs *WT* groups and *dec2*$^{P384R}$ vs *dec2*$^{WT}$ groups. GO terms and KEGG pathway with p-adj < 0.05 are significantly enriched.

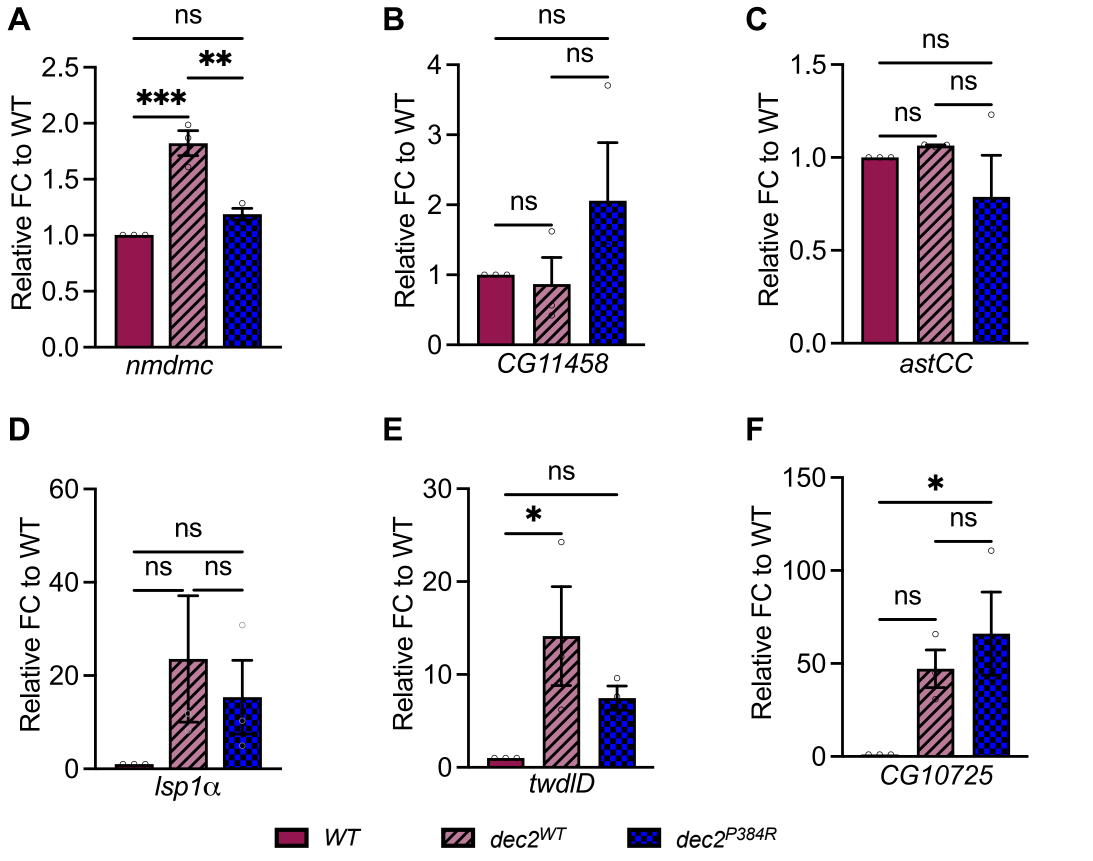


**Supplemental Figure 5: qPCR analysis of candidate genes from the RNAseq data. A-F.** Expression of *nmdmc* (A), *CG11458* (B), *astCC* (C), *Isp1α* (D), *twdlD* (E) and *CG10725* (F) measured by qPCR (n=3). FC=fold change, ns=not significant, *p<0.05, **p<0.01, ***p<0.001; one-way ANOVA with Tukey's comparisons.

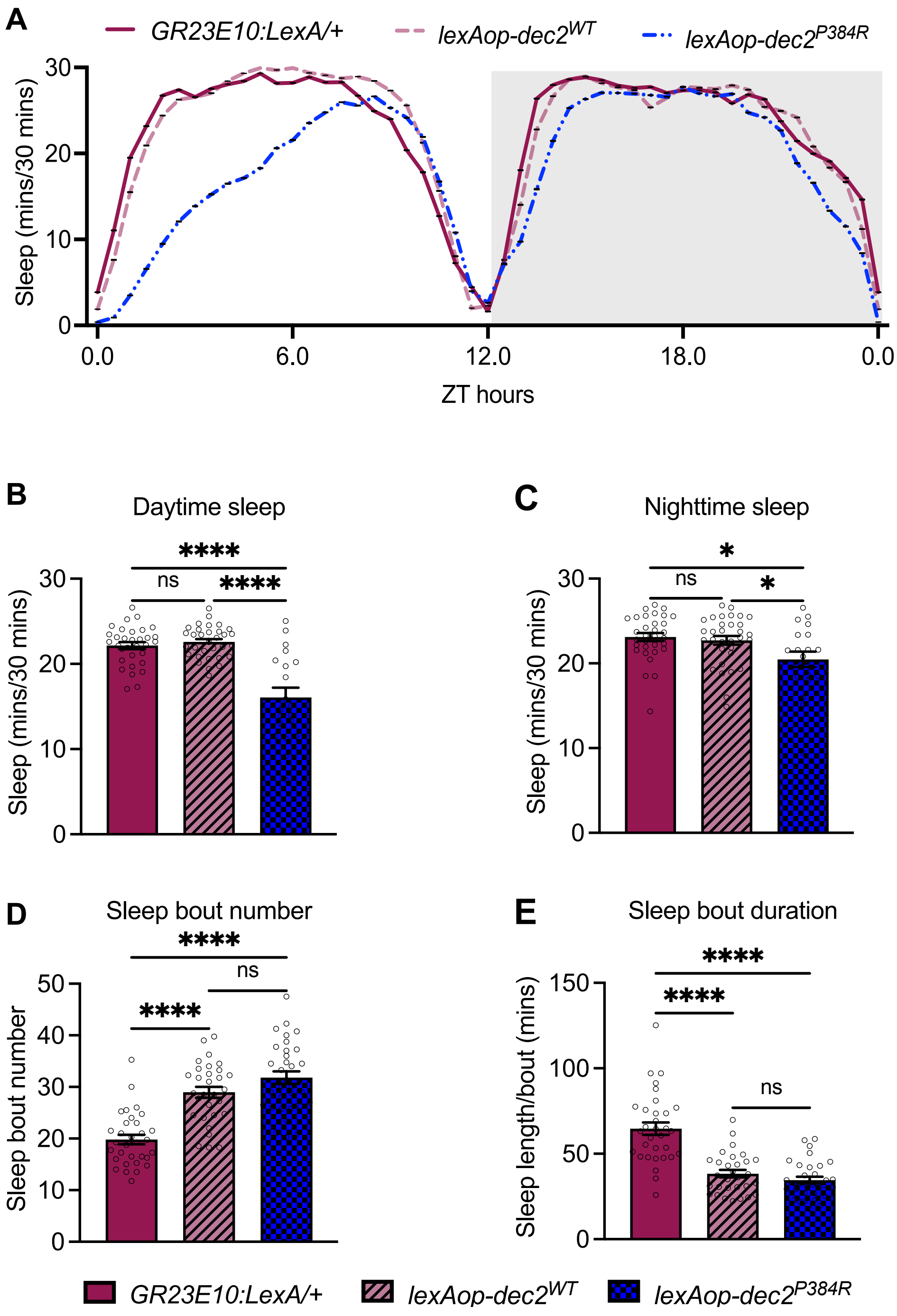


**Supplemental Figure 6: Sleep analysis of *dec2* transgenes expressed in sleep neurons with the lexA/lexAop expression system. A.** Sleep analysis in 12:12h L:D condition for the *GR23E10:LexA/+* (n=32)*, lexAop-dec2*$^{WT}$ (n=32) and *lexAop-dec2*$^{P384R}$ genotypes (n=20). **B-C.** Average sleep during daytime (B) and nighttime (C) of the genotypes indicated. **D-E.** Average sleep bout number (D) and sleep length/bout (E) in the genotypes indicated. ns=not significant, *p<0.05, ****p<0.0001; one-way ANOVA with Tukey's comparisons.

*mtnA*
Forward primer: CGGAAGCGGATGCAAATG
Reverse primer: CCGCAGGCGGATTTCTT
*mtnB*
Forward primer: GAACAAACTGCCAGTGCTC
Reverse primer: GGCCCATTCTTGCAAACG
*mtnC*
Forward primer: CAAAGGCTGCGGAACAAAC
Reverse primer: TGCTCTTGCAACACTGATCT
*mtnD*
Forward primer: ACCAAGTGCGGTGACAA
Reverse primer: GGAGCAGCACTTGTCCTT
*mtnE*
Forward primer: CAAGGGATGTGGAAACAACTG
Reverse primer: GCATTGGCATTGGCTGTTTC
*CG11699*
Forward primer: CCTTCTGCTACGGCTTCTG
Reverse primer: GGTTCGTCCTTGGTGATTCT
*mthl8*
Forward primer: TACGTGCAGTACGGACTTTC
Reverse primer: TGGACTAGATTTGGAGCAGTTT
*lsp1alpha*
Forward primer: TCTTCCCGCAGTTCTTCTTC
Reverse primer: GGCTAGTCTTCATCCACATCTC
*nmdmc*
Forward GAGCGTAATTGTGTCGAGGAA
Reverse GGCAGATCCAGCGAGTTATTAG
*CG11458*
Forward primer: ATGCGCTTCGCTCTACTT
Reverse: TCCTCCTCCTCCATTCTTCT
*CG10725*
Forward primer: GTGGGCAACTGTAGCAAGTA
Reverse primer: CAATACACTCGACCTCCATCAG
*twdlD*
Forward primer: GCCGATAAACTGGGCTACAA
Reverse primer: CTCCAATCACCTGACCACTTC
*astCC*
Forward primer: TCCGTTGTGCAGAACCTATC
Reverse primer: GCCCGGATACACCATCAAT

**Figure S7:** List of primers used for qPCR

**A**

| Descriptive statistics | | | |
|---|---|---|---|
| Genotype | n | Mean (days) | SEM |
| Figure 3A | | | |
| *WT* | 93 | 63.34 | 1.58 |
| *dec2* $^{WT}$ | 93 | 59.97 | 1.56 |
| *dec2* $^{P384R}$ | 66 | 81.65 | 2.60 |
| Figure 3B | | | |
| *WT* | 115 | 9.27 | 0.55 |
| *dec2* $^{WT}$ | 97 | 16.49 | 0.64 |
| *dec2* $^{P384R}$ | 85 | 15.59 | 0.71 |
| Figure 3C | | | |
| *WT* | 116 | 46.59 | 1.24 |
| *dec2* $^{WT}$ | 98 | 38.52 | 0.91 |
| *dec2* $^{P384R}$ | 104 | 59.75 | 1.11 |
| Figure 3D | | | |
| *WT* | 112 | 4.81 | 0.33 |
| *dec2* $^{WT}$ | 105 | 9.28 | 0.55 |
| *dec2* $^{P384R}$ | 128 | 8.23 | 0.34 |

| Genotype | n | Mean (hours) | SEM |
|---|---|---|---|
| Figure 4J | | | |
| *WT* | 185 | 51.50 | 0.46 |
| *dec2* $^{WT}$ | 129 | 52.56 | 0.63 |
| *dec2* $^{P384R}$ | 176 | 80.45 | 0.97 |

| Genotype | n | Mean (days) | SEM |
|---|---|---|---|
| Figure 4K | | | |
| *WT* | 96 | 6.20 | 0.13 |
| *dec2* $^{WT}$ | 94 | 5.51 | 0.15 |
| *dec2* $^{P384R}$ | 110 | 6.91 | 0.14 |

| Genotype | n | Mean (hours) | SEM |
|---|---|---|---|
| Figure 6 | | | |
| *GR23E10:LexA/+* | 43 | 39.67 | 1.03 |
| *lexAop-dec2* $^{WT}$ | 44 | 44.45 | 1.47 |
| *lexAop-dec2* $^{P384R}$ | 73 | 56.38 | 1.16 |
| *lexAop-dec2* $^{P384R}$*/mtnB* $^{i}$ | 70 | 44.23 | 0.92 |
| *lexAop-dec2* $^{P384R}$*/mtnC* $^{i}$ | 117 | 63.18 | 1.10 |
| *lexAop-dec2* $^{P384R}$*/mtnD* $^{i}$ | 111 | 55.68 | 0.94 |
| *lexAop-dec2* $^{P384R}$*/nmdmc* $^{i}$ | 54 | 59.11 | 1.36 |
| *lexAop-dec2* $^{P384R}$*/CG11699* $^{i}$ | 37 | 49.62 | 0.67 |

**B**

| Log-rank test results | | | |
|---|---|---|---|
| Condition | χ2 | p-value | Corrected p-value |
| Figure 3A | | | |
| *WT v.s. dec2* $^{WT}$ | 3.02 | 0.0823 | 0.1645 |
| *WT v.s. dec2* $^{P384R}$ | 70.11 | 0.00E+00 | 0.00E+00 |
| *dec2* $^{WT}$ *v.s. dec2* $^{P384R}$ | 76.88 | 0.00E+00 | 0.00E+00 |
| Figure 3B | | | |
| *WT v.s. dec2* $^{WT}$ | 75.19 | 0.00E+00 | 0.00E+00 |
| *WT v.s. dec2* $^{P384R}$ | 40.27 | 0.00E+00 | 0.00E+00 |
| *dec2* $^{WT}$ *v.s. dec2* $^{P384R}$ | 2.07 | 0.1502 | 0.3004 |
| Figure 3C | | | |
| *WT v.s. dec2* $^{WT}$ | 32.91 | 9.60E-09 | 1.90E-08 |
| *WT v.s. dec2* $^{P384R}$ | 42.52 | 0.00E+00 | 0.00E+00 |
| *dec2* $^{WT}$ *v.s. dec2* $^{P384R}$ | 151.56 | 0.00E+00 | 0.00E+00 |
| Figure 3D | | | |
| *WT v.s. dec2* $^{WT}$ | 49.45 | 0.00E+00 | 0.00E+00 |
| *WT v.s. dec2* $^{P384R}$ | 38.49 | 0.00E+00 | 0.00E+00 |
| *dec2* $^{WT}$ *v.s. dec2* $^{P384R}$ | 11.34 | 0.0008 | 0.0015 |

| Condition | χ2 | p-value | Corrected p-value |
|---|---|---|---|
| Figure 4J | | | |
| *WT v.s. dec2* $^{WT}$ | 2.33 | 0.1271 | 0.2543 |
| *WT v.s. dec2* $^{P384R}$ | 325.88 | 0.00E+00 | 0.00E+00 |
| *dec2* $^{WT}$ *v.s. dec2* $^{P384R}$ | 271.72 | 0.00E+00 | 0.00E+00 |

| Condition | χ2 | p-value | Corrected p-value |
|---|---|---|---|
| Figure 4K | | | |
| *WT v.s. dec2* $^{WT}$ | 8.82 | 0.003 | 0.006 |
| *WT v.s. dec2* $^{P384R}$ | 19.95 | 7.90E-06 | 1.60E-05 |
| *dec2* $^{WT}$ *v.s. dec2* $^{P384R}$ | 49.71 | 0.00E+00 | 0.00E+00 |

| Condition | χ2 | p-value | Corrected p-value |
|---|---|---|---|
| Figure 6 | | | |
| *GR23E10:LexA/+ v.s. GR23E10>lexAop-dec2WT* | 7.66 | 0.0057 | 0.017 |
| *GR23E10:LexA/+ v.s. GR23E10>lexAop-dec2* $^{P384R}$ | 65.31 | 0.00E+00 | 0.00E+00 |
| *GR23E10>lexAop-dec2* $^{P384R}$ *v.s. GR23E10>lexAop-dec2* $^{WT}$ | 34.51 | 4.20E-09 | 1.70E-08 |
| *GR23E10:LexA/+ v.s. lexAop-dec2* $^{P384R}$*/mtnB* $^{i}$ | 12.71 | 0.0004 | 0.0025 |
| *GR23E10:LexA/+ v.s. lexAop-dec2* $^{P384R}$*/mtnC* $^{i}$ | 117.04 | 0.00E+00 | 0.00E+00 |
| *GR23E10:LexA/+ v.s. lexAop-dec2* $^{P384R}$*/mtnD* $^{i}$ | 75.43 | 0.00E+00 | 0.00E+00 |
| *GR23E10:LexA/+ v.s. lexAop-dec2* $^{P384R}$*/nmdmc* $^{i}$ | 61.7 | 0.00E+00 | 0.00E+00 |
| *GR23E10:LexA/+ v.s. lexAop-dec2* $^{P384R}$*/CG11699* $^{i}$ | 36.27 | 0.00E+00 | 0.00E+00 |
| *GR23E10>lexAop-dec2* $^{P384R}$ *v.s. lexAop-dec2* $^{P384R}$*/mtnB* $^{i}$ | 57.87 | 0.00E+00 | 0.00E+00 |
| *GR23E10>lexAop-dec2* $^{P384R}$ *v.s. lexAop-dec2* $^{P384R}$*/mtnC* $^{i}$ | 17.49 | 2.90E-05 | 0.0001 |
| *GR23E10>lexAop-dec2* $^{P384R}$ *v.s. lexAop-dec2* $^{P384R}$*/mtnD* $^{i}$ | 0.3 | 0.5809 | 1 |
| *GR23E10>lexAop-dec2* $^{P384R}$ *v.s. lexAop-dec2* $^{P384R}$*/nmdmc* $^{i}$ | 2.55 | 0.11 | 0.7702 |
| *GR23E10>lexAop-dec2* $^{P384R}$ *v.s. lexAop-dec2* $^{P384R}$*/CG11699* | 20.44 | 6.20E-06 | 1.80E-05 |
| *GR23E10>lexAop-dec2* $^{WT}$ *v.s. lexAop-dec2* $^{P384R}$*/mtnB* $^{i}$ | 0.08 | 0.7836 | 1 |
| *GR23E10>lexAop-dec2* $^{WT}$ *v.s. lexAop-dec2* $^{P384R}$*/mtnC* $^{i}$ | 75.07 | 0.00E+00 | 0.00E+00 |
| *GR23E10>lexAop-dec2* $^{WT}$ *v.s. lexAop-dec2* $^{P384R}$*/mtnD* $^{i}$ | 35.48 | 2.60E-09 | 1.00E-08 |
| *GR23E10>lexAop-dec2* $^{WT}$ *v.s. lexAop-dec2* $^{P384R}$*/nmdmc* $^{i}$ | 36.94 | 0.00E+00 | 0.00E+00 |
| *GR23E10>lexAop-dec2* $^{WT}$ *v.s. lexAop-dec2* $^{P384R}$*/CG11699* $^{i}$ | 6.1 | 0.0135 | 0.0404 |

**Figure S8: A.** Descriptive statistics for lifespans in figure 3. **B.** Log-rant test results for lifespans.
